# Regulation of BCL11A DNA binding and expression in human erythrocyte precursor HUDEP-2 cells

**DOI:** 10.64898/2026.02.06.704516

**Authors:** Meigen Yu, Puspa Das, John R. Horton, Jujun Zhou, Jisun Lee, Tingting Hong, Yue Lu, Marcos R. Estecio, Polina A. Iakova, Abhinav K. Jain, Gianluca Sbardella, Yan Xiong, Jian Jin, Robert M. Blumenthal, Yun Huang, Xing Zhang, Xiaodong Cheng

**Author notes:** These authors contributed equally (Meigen Yu, Puspa Das).

## Abstract

BCL11A is a transcription factor crucial for neurodevelopment and hematopoiesis. It regulates the developmental switch from fetal hemoglobin (HbF) to adult hemoglobin and is a major therapeutic target for sickle cell disease and β-thalassemia. BCL11A exists in multiple isoforms, including the L isoform (containing a single two-finger ZF2-3 DNA-binding domain) and the XL isoform (containing two arrays: the two-finger ZF2-3 and the three-finger ZF4-6). We used three approaches to investigate BCL11A functions. First, we examined DNA recognition by BCL11A, which preferentially binds the short 6-bp DNA motif TGNCCA. ZF4-6 recognizes all four variants of this motif with distinct strand-specific interactions: TGTCCA on the top strand, TG(A/C)CCA on the complementary strand, and the palindromic TGGCCA on either strand. ZF2-3 also binds TGTCCA from the top strand, featuring a unique thymine interaction by ZF2 Phe388. Motif multiplicity within BCL11A binding sites may promote BCL11A oligomerization by enabling multiple ZF arrays to engage DNA simultaneously. Second, we treated HUDEP-2 cells (which express adult hemoglobin) with inhibitors targeting three epigenetic silencing marks **–** DNA methylation, histone H3 lysine 9 methylation or H3 lysine 27 methylation. All treatments, individually or in combination, increased HbF expression to varying degrees. Notably, FTX6058 markedly reduced BCL11A transcription and translation (likely via effects on LIN28B), while EML741 caused a partial reduction. Third, we screened 213 pomalidomide- and lenalidomide-derived compounds and quantified proportions of HbF+ cells by flow cytometry. Effects of four compounds were analyzed by protein mass spectrometry. Although BCL11A levels themselves were unchanged, all four compounds selectively decreased levels of known pomalidomide targets, with consistently decreased levels of the zinc-finger proteins IKZF1 and ZFP91. Together, our studies clarify how BCL11A recognizes DNA, how its expression can be modulated epigenetically, and how small-molecule degraders influence its regulatory network, providing new avenues for HbF reactivation therapies.

## INTRODUCTION

Both activation and repression of gene transcription rely on sequence-specific DNA binding by transcription factors^1–3^. In adult human erythroid cells, silencing of the *HBG1* and *HBG2* genes – which encode the γ-globin chains that pair with α-globin chains to form α_2_γ_2_ fetal hemoglobin (HbF) – is mediated directly or indirectly by several C2H2 zinc-finger proteins. These include <u>B</u>-<u>c</u>ell lymphoma/leukemia <u>11A</u> (BCL11A)^4^, <u>K</u>rüppel-like <u>f</u>actor <u>1</u> (KLF1)^5,6^, <u>Z</u>inc Finger and <u>BTB</u> Domain Containing <u>7A</u> (ZBTB7A; also known as leukemia/lymphoma-related factor, LRF)^7^, <u>Z</u>i<u>n</u>c <u>F</u>inger Protein <u>410</u> (ZNF410)^8,9^, and <u>H</u>ypermethylated in <u>C</u>ancer <u>2</u> (HIC2)^10^. Among these, erythroid lineage-specific disruption of *BCL11A* expression, using CRISPR-Cas9 editing^11^, formed the basis for one of the two gene therapies the FDA approved in 2023 for treating <u>s</u>ickle <u>c</u>ell <u>d</u>isease (SCD) ^footnote1^. In 2024, a 12-year-old boy became the first SCD patient to complete this treatment ^footnote2^.

However, the impact of gene therapies for SCD is likely to remain limited in the near future. With costs currently estimated in the millions of dollars per treatment, access will be restricted to a very small number of patients^12^. Considering that more than 100,000 individuals live with SCD in the United States alone – and millions more worldwide – the need for affordable and broadly effective therapies for hemoglobinopathies remains urgent. The vast majority of affected patients, even those with healthcare insurance, will not have access to gene therapy in the foreseeable future.

An alternative, generally less-expensive, approach to SCD therapy involves small molecule agents, although current options are suboptimal^13^. Here we investigate the ability of three small-molecule epigenetic inhibitors, used individually and in combination, to boost expression of HbF in <u>H</u>uman <u>U</u>mbilical cord blood-<u>D</u>erived <u>E</u>rythroid <u>P</u>rogenitor (HUDEP-2) cells^14,15^, which normally express adult hemoglobin (α_2_β_2_). These inhibitors are GSK3484862 – a DNMT1-specific DNA methyltransferase inhibitor^16^, EML741 – an inhibitor of euchromatic histone-lysine methyltransferases EHMT1/2 (G9a/GLP)^17^, and FTX6058 **–** a compound targeting the <u>e</u>mbryonic <u>e</u>ctoderm <u>d</u>evelopment (EED) subunit of <u>p</u>olycomb <u>r</u>epressive <u>c</u>omplex <u>2</u> (PRC2)^18^. These agents act, respectively, by reducing three repressive chromatin marks: DNA CpG methylation, histone H3 lysine 9 dimethylation (H3K9me2), and H3 lysine 27 trimethylation (H3K27me3).

Previous studies support the therapeutic potential of these agents. In a transgenic mouse model of SCD, oral administration of GSK3482364 was well tolerated and increased both overall HbF levels and the proportion of erythrocytes expressing HbF^19^. The GSK3484862 used in our study is the purified *R*-enantiomer of GSK3482364 that selectively targets DNMT1 for degradation^20^. Inhibition of G9a with RK-701 reduces H3K9 methylation and boosts levels of <u>b</u>eta-<u>g</u>lobin locus transcript <u>3</u> (*BGLT3*)^21^, which is a long non-coding RNA coded upstream of the *HBBP1* pseudogene, which in turn is implicated in HbF induction^22,23^. The G9a inhibitor used here, EML741, also strongly inhibits G9a while exhibiting improved selectivity, low cell toxicity, and favorable permeability relative to other G9a inhibitors^17^. Finally, FTX6058 has been reported to boost HbF levels, though to date only in multiple meeting abstracts ^footnote3^ as well as in a completed Phase 1 clinical trial report (NCT04586985).

We also revisited pomalidomide-mediated HbF induction^24,25^. Pomalidomide and its analog lenalidomide bind to cereblon, a component of an E3 ubiquitin ligase complex, and trigger the degradation of IKZF1 and IKZF3 in multiple myeloma cells^26,27^. However, genetic ablation of IKZF1 does not recapitulate the effect of pomalidomide treatment in SCD-derived CD34^+^ <u>h</u>ematopoietic <u>s</u>tem and <u>p</u>rogenitor <u>c</u>ells (HSPCs)^28^, suggesting that additional or alternative targets may underlie this HbF induction. Notably, we noticed that ZNF410 shares 50% sequence identity in two of its zinc fingers with IKZF1 ZF2 – the primary pomalidomide-binding site in IKZF1 – and that ZF5 of ZNF410 harbors the key residues required for pomalidomide binding^29^. These sequence similarities led us to screen more than 200 pomalidomide derivatives for additional targets in HUDEP-2 cells.

In addition to targeting the *HBG1* and *HBG2* genes, BCL11A binds thousands of other genomic sites in HUDEP-2 cells. This was demonstrated by two independent experiments: (i) chromatin immunoprecipitation sequencing (ChIP-seq) of endogenously inducible BCL11A-ER-V5 using an anti-V5 antibody, which revealed the consensus binding motif TGNCCA (N=any nucleotide)^30^, and (ii) BCL11A CUT&RUN, which identified the related motif TGNCCW (W=A or T)^31^ (Figure 1A). Functional studies further showed that depletion of a BCL11A-FKBP12(K36V) fusion protein expressed in HUDEP-2 cells boosted expression of 44 genes^32^. However, as noted in Mehta et al.^32^, only half of these genes (22/44) contained the preferred BCL11A binding motif TGACCA in their promoters. Building on our recent structural characterization of the BCL11A DNA-binding ZF arrays^33^, we now demonstrate that the ZF4-6 three-finger array recognizes all four variants of the TGNCCA motif, but significantly does so through two distinct binding modes. In addition, for the first time, we determined the structures of the ZF2-3 two-finger array bound to DNA.

**Figure 1.**
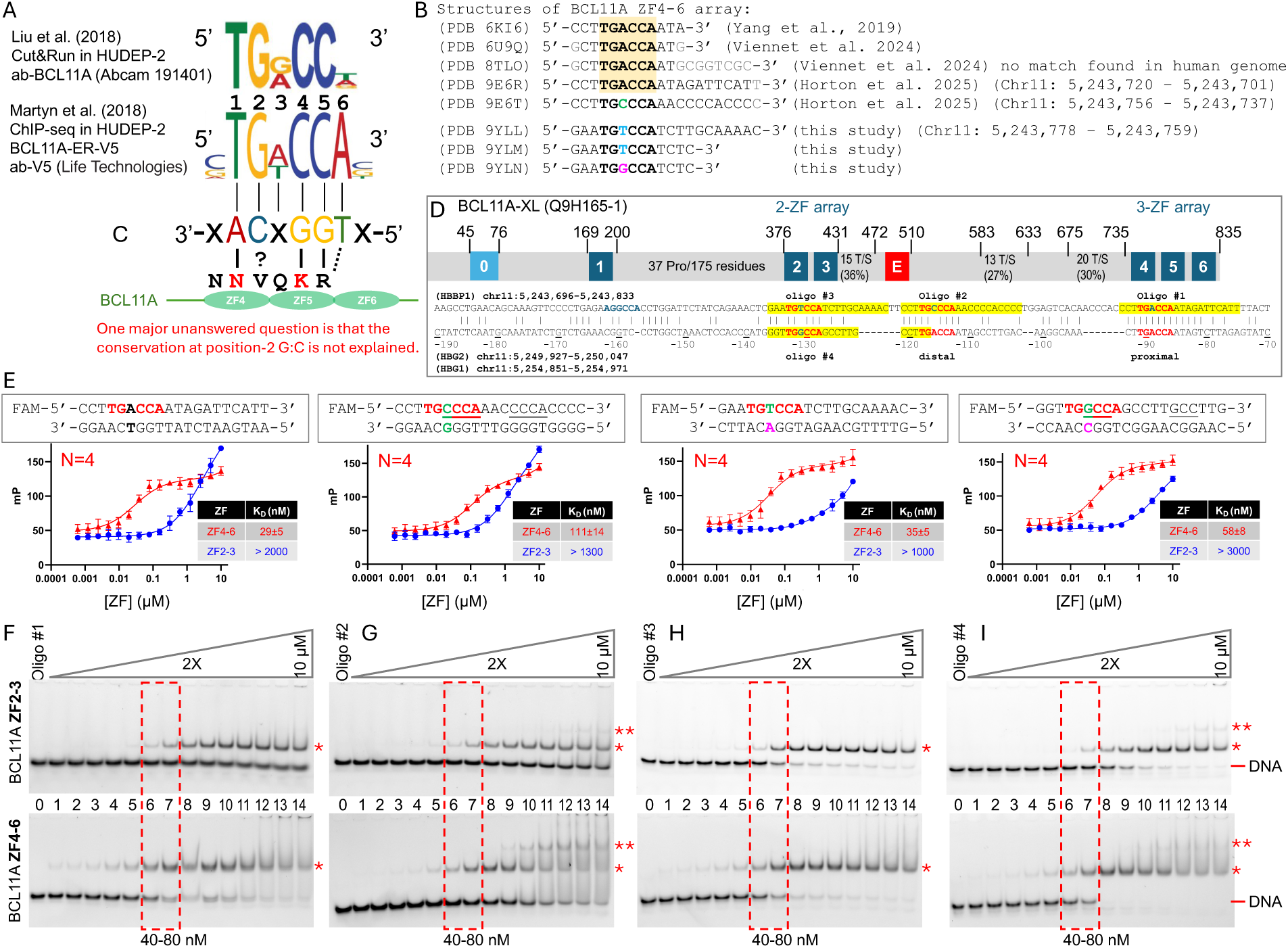
BCL11A ZF2-3 and ZF4-6 fragments bind all four variants of the TGNCCA motif. (**A**) Consensus of binding by BCL11A derived from CUT&RUN^31^ and ChIP-seq^30^ experiments in HUDEP-2 erythroid cells. Notable similarities include DNA sequence positions 1, 2, 4 and 5, but there are differences at positions 3 and 6. (**B**) List of available cocrystal structures of BCL11A ZF4-6 and the DNA sequences used. The common TGACCA sequences are shaded. (**C**) A cartoon shows the bottom-strand recognition mode by noncontiguous residues in ZF4-6. Solid lines indicate direct hydrogen bonds and dashed line indicates van der Waals contact. (**D**) The largest splicing isoform of BCL11A-XL and its corresponding UniProt accession number are shown. Each C2H2-ZF unit is numbered and highlighted in blue. The red box labeled ‘E’ indicates a tract of 20 consecutive acidic residues (mostly Glu) that may act in *cis* as a DNA mimic. The BCL11A binding site at the regulatory region between *BGLT3* and *HBBP1* contains three copies of TGNCCA (in red), aligned with the well-studied promoter regions of the two γ-globin genes (*HBG2* and *HBG1*). Four 20-bp oligos, #1, #2 and #3 of *HBBP1* (highlighted in yellow) and #4 of *HBG2/1* were synthesized with FAM labels for binding assays. (**E**) FP assays of ZF2-3 (blue lines) and ZF4-6 (red lines) arrays for binding four oligos with sequences indicated above. The number of replicates is indicated by N and error bars represent SEM. For binding curves of ZF2-3 (blue lines) that did not reach saturation, the lower limit of the binding affinity was estimated. (**F-I**) EMSA of ZF2-3 (upper panels) and ZF4-6 (lower panels) binding with four oligos. The red asterisks indicate putative protein-DNA complexes. Dashed red boxes indicate the area of the apparent K_D_ for ZF4-6.

## Results

### Conservation of guanine G2 of the BCL11A binding motif remains unexplained

Given that the experimentally-determined BCL11A binding motif is relatively short (six nucleotides), with four or five positions being conserved (1, 2, 4, 5 and **–** possibly to a lesser extent **–** 6) (Figure 1A), its sequence specificity can in theory be explained by currently available structural data of the C-terminal three-finger array (ZF4-6) bound to TG<u>A</u>CCA and TG<u>C</u>CCA (Figure 1B)^33–35^. Because CUT&RUN analysis indicated the sixth position is partially degenerate (either A or T)^31^ (Figure 1A), we focused on the TGNCCA core motif to simplify comparison between the CUT&RUN and ChIP-seq datasets, as well as with our *in vitro* binding results.

DNA sequence-specific recognition occurs primarily through the opposite strand of the motif: adenine at position 1 (A1) interacts with Asn756 of ZF4, and guanines at positions 4 and 5 (G4 and G5) interact with Lys784 and Arg787 of ZF5, respectively (solid lines in Figure 1C). The pairings of Arg/Lys with guanine and Asn with adenine are a well-established mechanism of purine recognition^36–38^. In addition, preference for an A:T base pair at position 6 – which is a partially degenerate position in CUT&RUN (Figure 1A) **–** can be attributed to an interaction between Arg787 and the thymine (T6) methyl group in the TpG dinucleotide (dashed line in Figure 1C). The T6 methyl group makes van der Waals contacts with Arg787, which simultaneously engages G5 on the same strand – together forming what we term ‘methyl-Arg-G triads’^39,40^. By contrast, conservation of guanine at position 2 (G2) remains unexplained. The corresponding residue, Val759 in ZF4, lies too far from the base pair, and valine, being hydrophobic, typically does not contribute to base-specific recognition.

BCL11A contains two DNA-binding zinc-finger arrays, ZF2-3 and ZF4-6, separated by a 300-amino-acid linker enriched in serine and threonine residues and containing a polyglutamate stretch (red box labeled ‘E’ in Figure 1D). To investigate their binding properties, we purified recombinant proteins of isolated ZF2-3 and ZF4-6 fragments and performed *in vitro* DNA binding assays using four 20-bp DNA oligonucleotides (Figure 1E-I). These four DNA duplexes were derived from the BCL11A binding sites between *BGLT3* and *HBBP1,* as well as from the *HBG1/2* promoters (Figure 1D). Fluorescence polarization (FP) assay showed that ZF2-3 binds DNA more weakly than ZF4-6, with apparent dissociation constants (K_D_) in the range of 1-3 μM for ZF2-3 and 0.03-0.11 μM for ZF4-6 (Figure 1 E). Among the test oligonucleotides, ZF4-6 bound most strongly to oligo #1 (containing a TG<u>A</u>CCA motif) and oligo #3 (containing TG<u>T</u>CCA), followed by oligo #4 (TG<u>G</u>CCA) with ∼2-fold reduced affinity, and finally oligo #2 (TG<u>C</u>CCA), which exhibited ∼4-fold weaker binding (Figure 1E). Electrophoretic mobility shift assay (EMSA) revealed that for oligos #1 (TG<u>A</u>CCA) and #3 (TG<u>T</u>CCA), both arrays produced a single shifted band (Figure 1F and 1H). In contrast, binding to oligos #2 (TG<u>C</u>CCA) and #4 (TG<u>G</u>CCA) generated two distinct shifts: the first appearing at ∼40-80 nM protein concentration for ZF2-3 and at ≤20 nM for ZF4-6, followed by a second shift emerging at higher protein concentrations (Figure 1G and 1I). Notably, for the ZF4-6 interaction with oligo #2, we previously observed two protein molecules bound to the same DNA duplex - one at the 5’ TG<u>CCCA</u> site and the other at the 3’ half overlapping the <u>CCCA</u> sequence^33^.

### Distribution of TGNCCA motifs within BCL11A Binding Sites

We next examined the genome-wide contribution of TGNCCA motifs using BCL11A chromatin occupancy data from HUDEP-2 cells. We analyzed five datasets from two independent studies: two replicates of ChIP-seq^30^ and three replicates of BCL11A CUT&RUN^31^. We built first-order Markov models for each replicate to estimate the expected random occurrence of TGNCCA sequences conditioned on the dinucleotide frequencies within the BCL11A binding sites. Overall, the observed TGNCCA frequency was increased by an average of 23.5% relative to expectation (p < 1×10^-20;^ Figure 2A; Supplementary Table S1). However, enrichment varied substantially among individual motif variants. TGACCA and TGTCCA were the most strongly enriched, on average more than twofold over expected (2.6- and 2.4-fold, respectively; p < 1×10^-300^), whereas TGGCCA occurred at approximately the expected rate in ChIP-seq (Figure 2A) or slightly above expected in the CUT&RUN data; Supplementary Figure S1). TGCCCA was the only underrepresented variant, occurring ∼22% less often than expected in a random sequence (0.78-fold; p < 1×10^-42^). These enrichment patterns closely match motif discovery results from the ChIP-seq analysis by Martyn *et al*. (Figure 1A)^30^. Notably, the ranking of motif enrichment also mirror our ZF4-6 binding affinities (K_D_ values of 29 nM for TGACCA < 35 nM for TGTCCA < 58 nM for TGGCCA < 111 nM for TGCCCA; Figure 1E), suggesting that BCL11A binding sites in the HUDEP-2 cell are preferentially enriched for the motifs it binds most strongly, and/or that potentially-competing sites are selected against.

**Figure 2.**
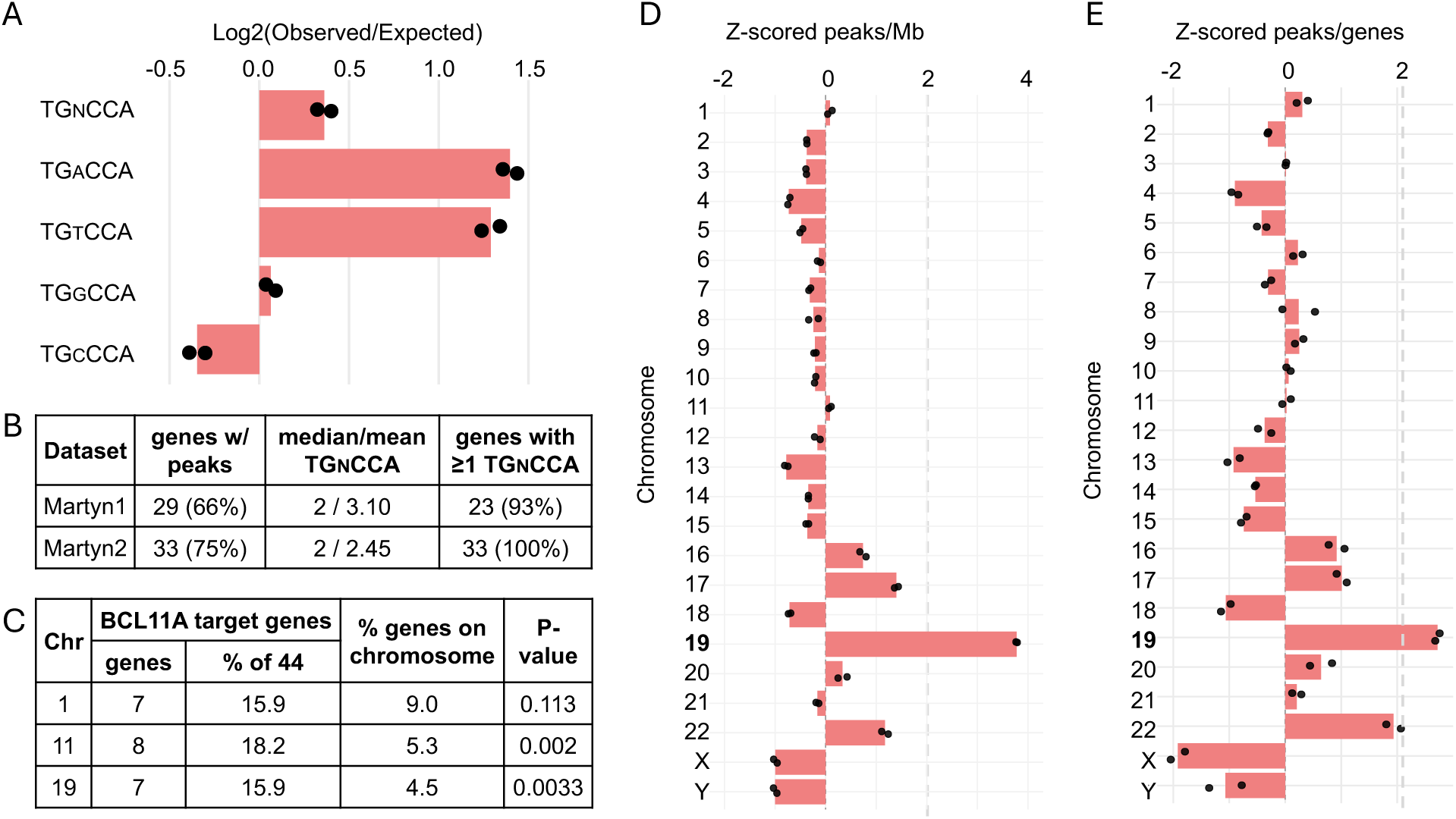
Distribution of BCL11A peaks and TGNCCA motifs in ChIP-seq data (hg38). (**A**) Enrichment of TGNCCA motifs in BCL11A ChIP-seq peaks, driven by higher-than-expected occurrence of TGACCA and TGTCCA. Expected motif occurrence was calculated using first-order Markov models for each replicate. (**B**) BCL11A ChIP-seq peaks associated with the 44 experimentally validated BCL11A target genes^32^, most of which contain multiple repeats of TGNCCA. (**C**) BCL11A target genes associated with BCL11A ChIP-seq peaks are enriched on chromosomes 11 and 19 by Fisher’s exact test. Columns (left to right) indicate chromosome number; the total number of genes associated with at least one BCL11A peak; the percentage of total 44 target genes represented; the percent of all annotated hg38 genes (both coding and non-coding) on that chromosome; and the corresponding Fisher’s exact test p-value. (**D, E**) Z-scored BCL11A peak density per megabase (Mb) or per hg38 annotated gene, showing strong enrichment on chromosome 19 and moderate enrichment on chromosomes 16, 17, and 22.

A previous study^32^ identified 44 genes repressed by BCL11A in HUDEP-2 cells, including 31 classified as primary targets (Figure 2B; Supplemental Table S2). Using HOMER^41^, we mapped BCL11A peaks associated with either the promoter or gene body of these genes and found that, for genes with associated peaks, practically all were associated with at least one TGNCCA motif. In addition, we observed a mean of 2-3 TGNCCA motifs per peak (Figure 2B; for examples see Figure 1D), suggesting that motif multiplicity may contribute to BCL11A oligomerization at these loci^33,42^. These BCl11A targets were unevenly distributed across the genome, with chromosomes 1, 11, and 19 each harboring 7 or 8 genes (Figure 2C), whereas most other chromosomes contained only 0-3 (Supplementary Table S3). Notably, only 6 of the 44 BCL11A target genes in HUDEP-2 cells overlapped with BCL11A targets identified in CD34+ HSPCs^43^, including the globin genes *HBG1/2* and *HBZ*. This limited overlap highlights both shared globin regulatory function and additional cell type-specific roles for BCL11A.

To assess chromosomal enrichment, we performed Fisher’s exact tests comparing the observed and expected number of BCL11A target genes (Figure 2C). Although chromosome 1 contained 7 of the 44 targets (16%), this did not represent significant enrichment (p = 0.113 OR = 1.9) given its large genomic gene content (9.0%). In contrast, chromosome 11 harbored 8 targets (18%) despite containing only 5.3% of annotated genes, representing a significant enrichment (p = 0.002, OR = 3.98). This enrichment may be driven in part by genes within the β-globin locus (*HBG1, HBG2,* and *HBBP1*). Similarly, chromosome 19 contained 7 target genes (16%), a significant enrichment given its 4.5% share of genes genome-wide (p = 0.0033, OR = 4.01). We next examined the chromosomal distribution of BCL11A binding peaks. When normalized by either chromosome length or gene count, chromosome 19 showed the strongest enrichment, with a z-score > 2 (Figure 2D–2E). Chromosomes 16, 17, and 22 also displayed above-average peak densities, though to a lesser extent. Notably, these four chromosomes, and chromosome 19 in particular, are among the most gene dense in the genome and exhibit high chromatin accessibility, two strongly correlated features^44^ that may contribute to the observed enrichment.

### An alternative DNA binding mode of ZF4-6

Recently we structurally characterized ZF4-6 bound to oligo #1 (TG<u>A</u>CCA) or oligo #2 (TG<u>C</u>CCA)^33^. In both instances, ZF4-6 recognizes the complementary strand of the sequence motif and anchors the protein through three sequence-specific interactions: adenine with Asn756 of ZF4, and guanines with Lys784 and Arg787 of ZF5 (Figure 3A). Unexpectedly, when ZF4-6 was bound to oligo #3 (TG<u>T</u>CCA), the protein engages the top strand instead (Figure 3B-C). Consistent with DNA binding assays, only one ZF4-6 molecule bound the 20-bp oligo #3, leaving the 3’ half of the DNA unoccupied. We shortened the DNA length to a 12-bp duplex with a 3’-overhang on both strands (G or C in Figure 3D), which yielded a higher-resolution structure at 1.9 Å (Figure 3E). Both structures revealed the same DNA binding mode, with ZF4-6 occupying the top strand. The protein orientation was unambiguously determined by the anomalous electron density of the zinc atoms (Figure 3F).

**Figure 3.**
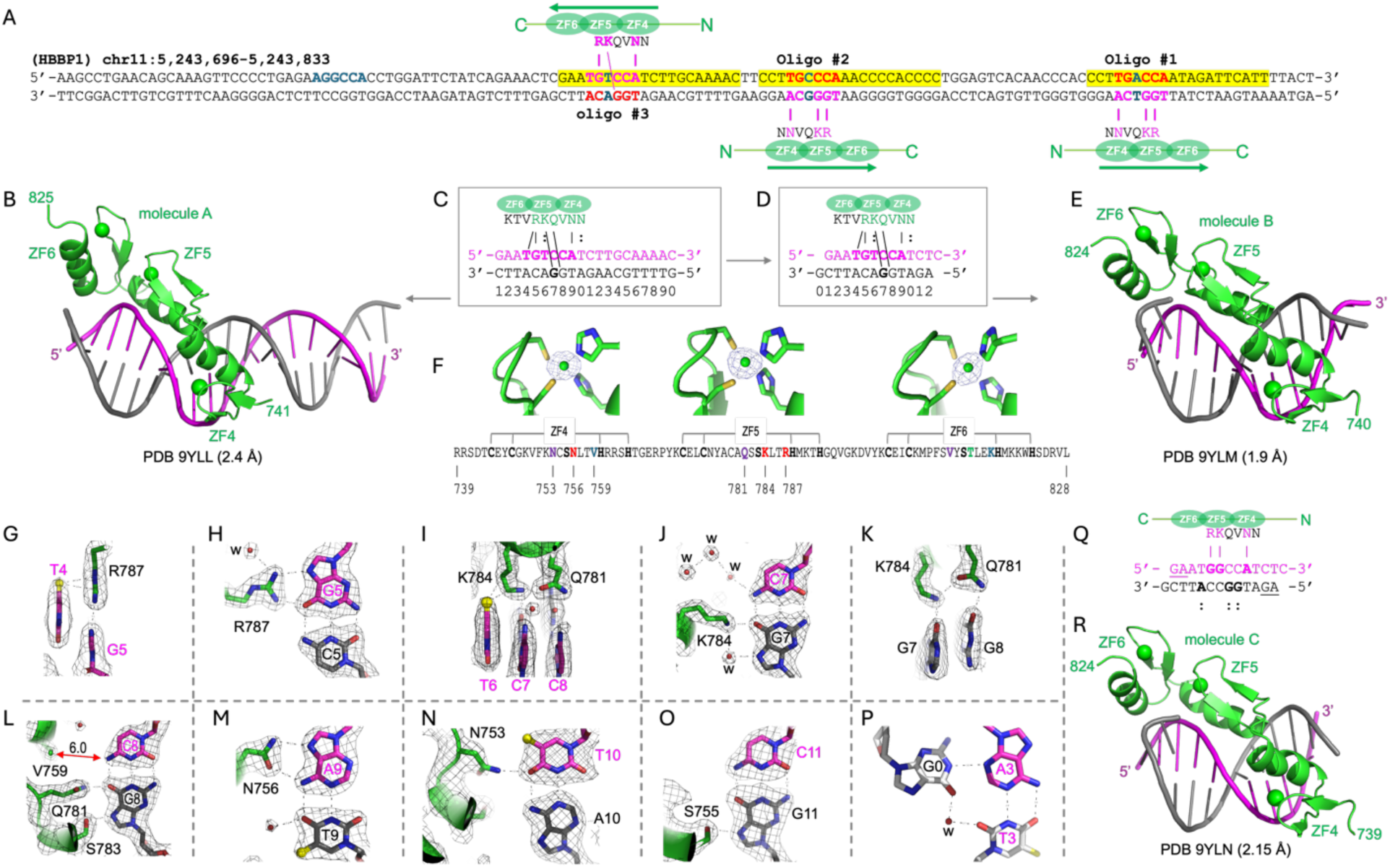
Structures of ZF4-6 bound with DNA in the top-strand recognition mode. (**A**) *HBBP1* regulatory region contains three variants of TGNCCA: TG<u>A</u>CCA (oligo #1), TG<u>C</u>CCA (oligo #2), and TG<u>T</u>CCA (oligo #3). These are recognized by ZF4-6 in either the top-strand mode (oligo #3) or bottom-strand mode (oligos #1 and #2). The arrows indicate the direction from amino to carboxy termini. (**B-C**) Structure of ZF4-6 bound with a 20-bp DNA. (**D-E**) Structure of ZF4-6 bound with a 12-bp DNA with 3’ overhangs. Green spheres represent Zn atoms. (**F**) Protein sequence of ZF4-6 used in the study. Residues located at base-interacting positions within ZF4 and ZF5 are labeled. The corresponding anomalous electron densities for zinc atoms are contoured at 5.0σ above the mean in PDB 9YLM. (**G-P**) Detailed interactions in PDB 9YLM. The electron densities of 2Fo-Fc in grey meshes are contoured at 1.5σ above the mean. (**G**) Arg787 interacts with T4 and G5. (**H**) Arg787 interacts with G5. (**I**) Lys784 and Gln781 interact with three base pairs (positions 6-7). (**J**) Lys784 interacts with G7. (**K**) Lys784 and Gln781 intramolecularly interact. (**L**) Gln781 and Ser781 interact with G8. (**M**) Asn756 interacts with A9. (**N**) Asn753 interacts with T10. (**O**) Ser755 interacts with G11. (**P**) The 3’ overhang guanine (G0) contacts with A:T base pair at position 3 in the DNA minor groove. (**Q-R**) ZF4-6 bound with the fully palindromic sequence TG<u>G</u>CCA.

In the top strand binding mode, Arg787 of ZF5 recognizes the TpG at positions 4 and 5 of the crystallized DNA sequence (Figure 3D, 3G-H). Notably, the Arg787-G5 interaction (Figure 3H) provides the missing explanation for guanine recognition at position 2 of the T**<u>G</u>**NCCA motif, as discussed earlier. Lys784 and Gln781 of ZF5 span across the next three base pairs (6, 7 and 8 in Figure 3I). Lys784 makes van der Waals contact with T6 (Figure 3I), while its terminal amino group forms a cross-strand hydrogen bond with G7 of the opposite strand (Figure 3J). Similarly, Gln781 forms a cross-strand hydrogen bond with G8 and an intramolecule interaction with Lys784 (Figure 3K). In addition, Ser783 of ZF5 contributes to DNA binding by forming an additional hydrogen bond with G8 (Figure 3L).

For ZF4, Val759 is positioned too far away from the nearest DNA base (6.0 Å from C8) to contribute to base-specific recognition (Figure 3L). Asn756 establishes classical bidentate hydrogen bonds with A9 (Figure 3M), whereas Asn753 forms a hydrogen bond with T10 (Figure 3N). Similar to Ser783 of ZF5, Ser755 of ZF4 also supports DNA binding by forming a hydrogen bond with G11 (Figure 3O). Additionally, the 3’ overhang cytosine of the top strand is poorly ordered, while the 3’ overhang guanine of the bottom strand folds back into the minor groove and interacts with the A:T base pair at position 3 (Figure 3P).

Comparing the two binding modes – top-strand recognition of TGTCCA versus bottom-strand recognition of TGCCCA and TGACCA (Figure 3A) – the key difference lies in Lys784. This lysine residue interacts with the same guanine paired with cytosine at position 4 of the TGN**C**CA motif, but the interaction occurs either through a cross-strand contact in the top-strand mode or from the same strand in the bottom strand mode. Because the TGN**C**CA motif is pseudo-symmetric, with outer T:A and G:C base pairs, the Asn756-A (positions 1 or 6) and Arg787-G (position 2 or 5) interactions are conserved in both modes.

Next, we substituted TG<u>T</u>CCA with the fully palindromic sequence TG<u>G</u>CCA in the same 12-bp oligo context (Figure 3Q), in order to test the mode of ZF4-6 interaction with a symmetric sequence. Although ZF4-6 still bound the top strand – likely because the 3’-overhang G favored crystal packing – the binding of Asn756, Lys784, and Arg787 mirrored that of the bottom-strand recognition mode. In this configuration, they interacted with the respective adenine and guanines from the same strand (Figure 3R).

### Structure of ZF2-3 bound with DNA

While the ZF4-6 fragment crystallized readily with all four TGNCCA variants in both long (20-bp) and short (12-bp) DNA lengths, crystallization of ZF2-3 with DNA proved more challenging. After screening >20 DNA sequences of varying lengths, we successfully crystallized the ZF2-3 fragment with one TG<u>T</u>CCA element in a 12-bp blunt-end duplex (Figure 4A), obtaining crystals in two different crystallographic space groups (Figure 4B-C). Interestingly, similar to ZF4-6, ZF2-3 binds TG<u>T</u>CCA in the top-strand recognition mode. Notably, ZF2-3 and ZF4-5 share 73% sequence identity, including conservation of all four base-interacting residues between ZF3 (Q416, S418, K419, and R422) and ZF5 (Q781, S783, K784, and R787), and three out of four base-interacting residues between ZF2 (F388, S390, N391, and V394) and ZF4 (N753, S755, N756, V759) (Figure 4D), and these residues are all highly conserved evolutionarily.

**Figure 4.**
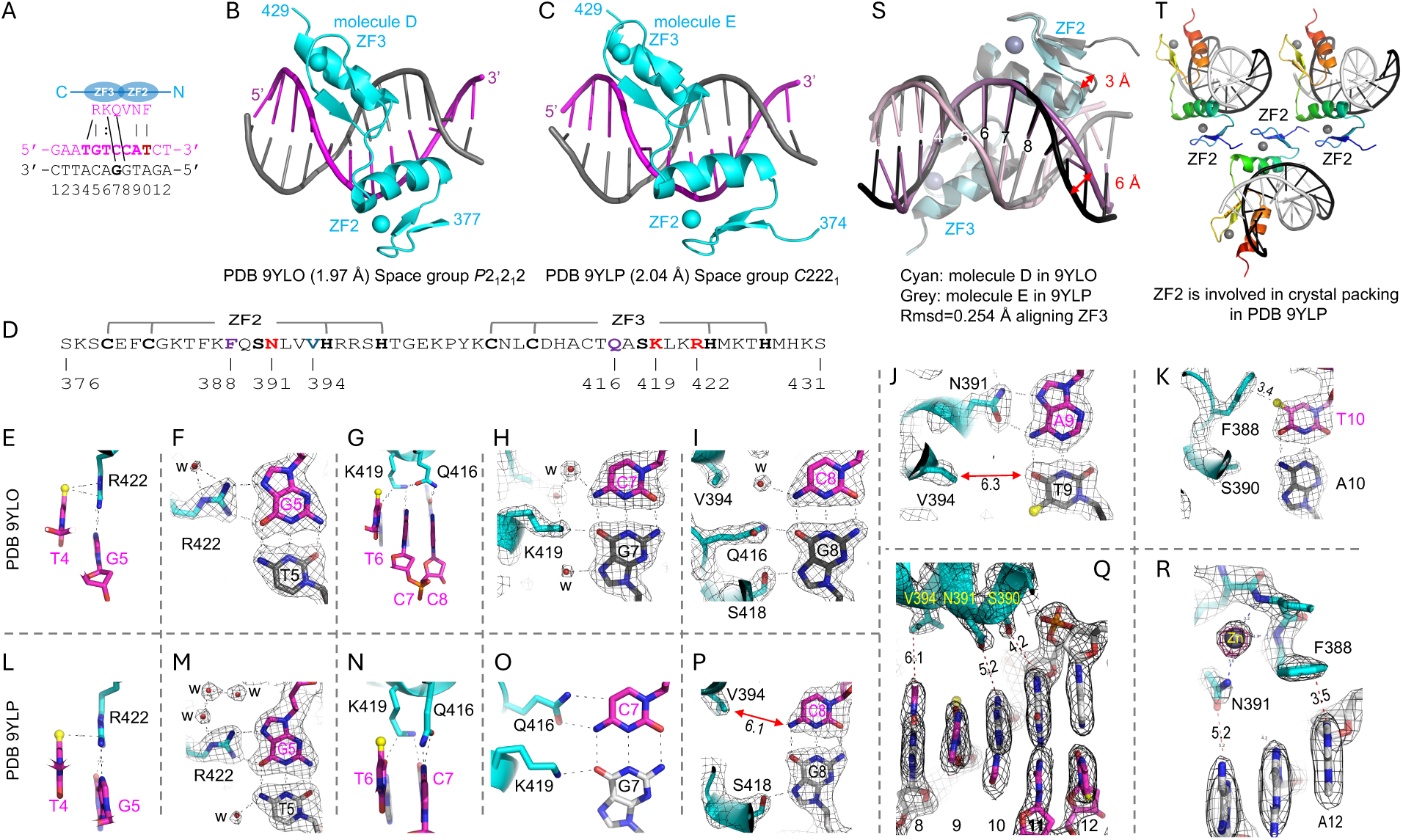
Structures of BCL11A ZF2-3 bound with DNA. (**A**) A 12-bp DNA was successfully crystalized with ZF2-3. (**B**) ZF2-3 complexed with DNA in space group *P*2_1_2_1_2 (PDB 9YLO). Spheres represent Zn atoms. (**C**) ZF2-3 complexed with DNA in space group *C*222_1_ (PDB 9YLP). (**D**) Protein sequence of ZF2-3 used in the study. Residues located at base-interacting positions within each finger are labeled. (**E-K**) Detailed interactions in space group *P*2_1_2_1_2 (PDB 9YLO). The electron densities of 2Fo-Fc in grey meshes are contoured at 1.5σ above the mean. (**E**) Arg422 interacts with T4 and G5. (**F**) Arg422 interacts with G5. (**G**) Lys419 and Gln416 interact with three base pairs (positions 6-8). (**H**) Lys419 interacts with G7. (**I**) Gln416 and Ser418 interact with G8. (**J**) Asn391 interacts with A9. (**K**) Phe388 interacts with T10. (**L-R**) Detailed interactions in space group *C*222_1_ (PDB 9YLP). The electron densities of 2Fo-Fc in grey meshes are contoured at 1.5σ above the mean. (**L**) Arg422 interacts with T4 and G5. (**M**) Arg422 interacts with G5. (**N**) Lys419 and Gln416 interact with two base pairs (positions 6-7). (**O**) Gln416 and Lys419 interact with C:G base pair at position 7. (**P**) Ser418 interacts with G8. (**Q**) Ser390, Asn391, and Val394 shift away from the DNA. (**R**) Phe388 shifted to interact with A12, from T10 (comparing to panel K). The anomalous electron density in orange meshes, contoured at 5.0σ above the mean, suggests a metal ion coordinated by side chain of Asn391 and main amide nitrogen atom of Phe388. (**S**) Superimposition of ZF2-3 bound with DNA in the two space groups. (**T**) ZF2 participates in crystal packing interactions in the *C*222_1_ space group (PDB 9YLP).

In the *P*2_1_2_1_2 structure (PDB 9YLO), ZF3 interacts with bases T_4_G_5_T_6_C_7_C_8_ in exactly the same manner as ZF5: Arg422 recognizes T4 and G5 (Figure 4E-F), Lys419 engages T6 and G7 (Figure 4G-H), and Gln416 together with Ser418 interacts with G8 (Figure 4I). Similarly to Val759 of ZF4, Val394 of ZF2 is positioned too far away to make direct base contacts, while Asn391 of ZF2 interacts with A9 (Figure 4J). A unique interaction arises from Phe388 of ZF2, whose aromatic ring stacks against the methyl group of T_10_, forming a CH-π interaction (Figure 4K). This distinct contact may explain the successful crystallization of ZF2-3, as a 3’ thymine (T10) following the TGNCCA motif was the only sequence context that yielded crystals. Aromatic residues are not commonly employed at base-interacting positions; however, ZF1 of ZNF410 utilizes two aromatic residues (Tyr and Trp) to interact with the methyl groups of two thymine bases^8^.

In the *C*222_1_ structure (PDB 9YLP), ZF3 remains tightly engaged with DNA (Figure 4L-O), while ZF2 shifts away from DNA, increasing the distance between its key residues (Val394, Asn391, Ser390, and Phe388) and their corresponding bases (Figure 4P-R). This difference is evident when the two structures are superimposed: ZF3 and its associated DNA base pairs (positions 4 to 8) overlap closely, with a root-mean-squared deviation of 0.254 Å, while ZF2 and its associated DNA base pairs (positions 9-12) are displaced by 3-6 Å (Figure 4S). We reasoned that this displacement is partly due to ZF2 participating in crystal packing interactions with ZF2 from neighboring molecules in the *C*222_1_ space group (Figure 4T).

### Inhibiting epigenetic silencing marks boosts HbF expression in HUDEP-2 cells

We next tested the effects of three epigenetic inhibitors, GSK3484862 (abbreviated as GSK), EML741 (EML), and FTX6058 (FTX) in HUDEP-2 cells. We followed a previously established three-phase liquid culture protocol (expansion, differentiation-growth, and differentiation-maturation) over 9-days for culturing HUDEP-2 cells^15^, administering compounds three times at media changes on day 0, day 3, and day 7 (Figure 5A). Cells were harvested at the ends of each phase (stage 1 at day 3, stage 2 at day 7, or stage 3 at day 9) for different experiments. First, we used western blots to test whether each of the three inhibitors effectively engaged their targets after 3 days of treatment in stage 1. GSK (DNMT1-inhibitor) induced DNMT1 degradation without any effect on cell viability (Figure 5B), consistent with prior observations in cancer cells ^20^. EML (G9a-inhibitor) reduced H3K9me2 levels, and FTX (EED-inhibitor) decreased H3K27me3 levels, as expected, again without impact on cell viability (Figure 5C-D). These reductions in DNMT1 and histone methylation marks were dose-dependent, leading us to select 0.1 μM FTX, 1 μM GSK, and 2 μM EML for further experiments. Second, we confirmed that compound treatment did not alter adult β-globin protein levels in stage 3 HUDEP-2 cells (Figure 5E-F). In contrast, we observed significant induction of fetal γ-globin following treatment with GSK and FTX (Figure 5E-F), but not with EML, even at concentrations up to 3 μM (Figure 5G). Prolonged inhibitor treatment did not significantly affect cell viability, as measured by Bio-Rad cell counter (Supplementary Figure S2A). Unexpectedly, FTX treatment markedly reduced BCL11A protein levels, while EML caused a partial reduction (Figure 5E, 5H).

**Figure 5.**
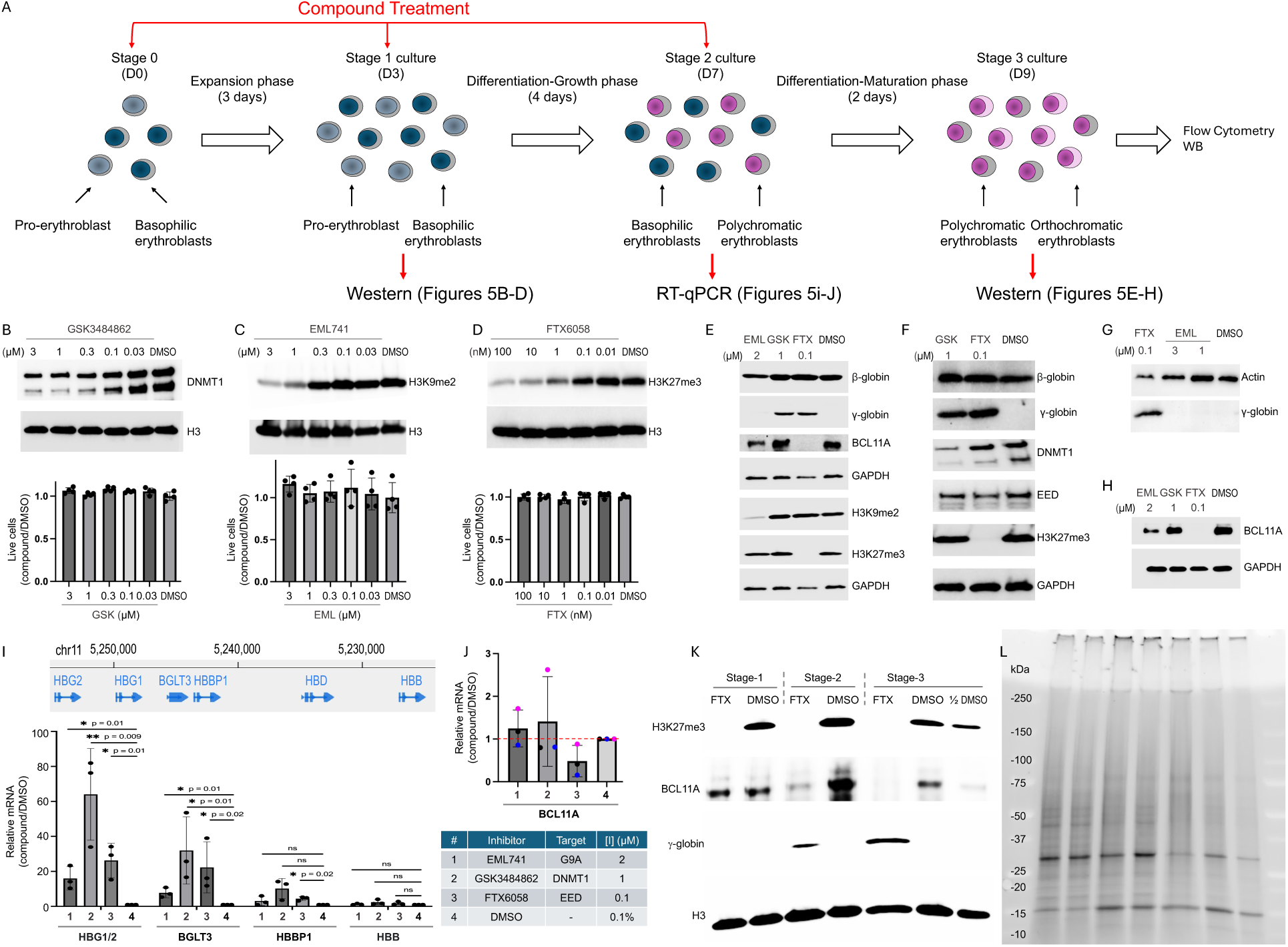
Inhibition of epigenetic silencing marks induces fetal globin expression in HUDEP-2 cells. (**A**) Schematic illustration of the HUDEP-2 cell culture differentiation workflow, showing compound treatment at day 0 (stage 0), day 3 (stage 1), and day 7 (stage 2). Cells at day 3 (stage 1), day 7 (stage 2), or day 9 (stage 3) were harvested for analysis. (**B-D**) HUDEP-2 cells were harvested at stage 1 after treatment with increasing concentrations of (**B**) GSK3484862 (GSK), (**C**) EML741 (EML), or (**D**) FTX6058 (FTX) for 3 days. Lower panels show the percentage of live cells quantified using a Bio-Rad cell counter and normalized to the DMSO control. Data represent mean ± SD from N=4 biological replicates. (**E-H**) HUDEP-2 cells were treated with each inhibitor (2 μM EML, 1 μM GSK, and 0.1 μM FTX). After 9 days of treatment, cells at stage 3 were harvested for western blot analysis. (**I**) Partial β-globin locus in chromosome 11 is shown above the relative mRNA expression levels (qPCR) in stage 2 HUDEP-2 cells. IP08 (protein transporter Importin 8) is used as a reference housekeeping gene. Data represent mean ± SD from N=3 independent experiments. **(J)** Relative mRNA expression levels of BCL11A. (**K-L**) Western blots and corresponding protein gel showing changes of H3K27me3 and BCl11A of HUDEP-2 cells at stages 1, 2 and 3.

We further analyzed transcript expression in stage 2 cells. Adult β-globin mRNA levels showed minimal changes (<2-fold increase) following compound treatments (Figure 5I), whereas fetal γ-globin mRNA increased by >20-60 fold. This induction appeared to correlate with elevated expression of *BGLT3*^21^ and *HBBP1*^45^ (Figure 5I). Notably, *BCL11A* expression at stage 2 was reduced by approximately 4-fold in FTX-treated cells (Figure 5J), following the reduction of the H3K27me3 mark observed at stage 1 (Figure 5K).

### FTX6058 activates RNA-binding protein LIN28B in HUDEP-2 cells

There are two known connections between BCL11A and the H3K27me3-catalyzing PRC2 complex - one at the protein level and the other at the mRNA level. In triple-negative breast cancer cells, BCL11A (via its N-terminal residues) directly interacts with PRC2 subunit RBBP4/7^46^. Whether a similar interaction occurs in HUDEP-2 cells remains to be determined. In CD34+ HSPC-derived proerythroblasts, treatment with the EZH2 enzymatic inhibitor EPZ-6438 activates the RNA-binding proteins IGF2BP1, IGF2BP3, and/or LIN28B^47^. LIN28B, in particular, regulates translation of *BCL11A* mRNA^48,49^. IGF2BP1 overexpression in cultured human adult erythroblasts induces fetal-like hemoglobin expression^50^ and is a key regulator of the γ-globin switch in K562 cells^51^. IGF2BP1 may have dual functions: it can bind *HBG1/2* mRNA to promote their translation, and it can bind and activate HIC2^52^, which itself represses BCL11A^10^. Taken together, a possible model is that the EED inhibitor FTX activated RNA-binding proteins modulate *BCL11A* transcription and translation, directly via LIN28B and indirectly via IGF2BP1.

In our experiments, LIN28B transcript expression was elevated in HUDEP-2 cells following 0.1 µM FTX treatment, whereas IGF2BP1 and HIC2 expression remained unchanged (Figure 6A). ChIP-qPCR analysis of histone modifications (active H3K4me3 and repressive H3K27me3) was performed at six gene loci (*LIN28B*, *IGF2BP1, HIC2*, *BCL11A* and *HBG1/2*). Upon FTX treatment, significant reductions in the repressive mark H3K27me3 were observed in *LIN28B*, *IGF2BP1, HIC2*, and *BCL11A* loci (Figure 6B). However, only the *LIN28B* locus exhibited a concomitant and significant increase in the activating mark H3K4me3 (Figure 6C), consistent with transcriptional activation. In contrast, no clear correlation between H3K4me3 and H3K27me3 levels was observed at *IGF2BP1* locus (Figure 6B-C). Both the *HIC2* and *BCL11A* loci contain distinct regions enriched for the activating acetylation (marked by H3K27ac) as well as regions marked by the repressive modification (H3K27me3), consistent with a bivalent chromatin feature (Supplementary Figure S3). In the repressive regions, FTX treatment led to a reduction of H3K27me3 accompanied by a small increase in H3K4me3, whereas in the regions of active chromatin, H3K4me3 levels were significantly reduced. We did not observe any notable changes in H3K4me3 or H3K27me3 at the promoters of *HBG1* and *HBG2* following FTX treatment. Together, these results suggest that, among the limited set of loci examined, FTX selectively remodels chromatin at the LIN28B locus by relieving the repressive H3K27me3 histone mark and promoting the activating H3K4me3 histone modification.

**Figure 6.**
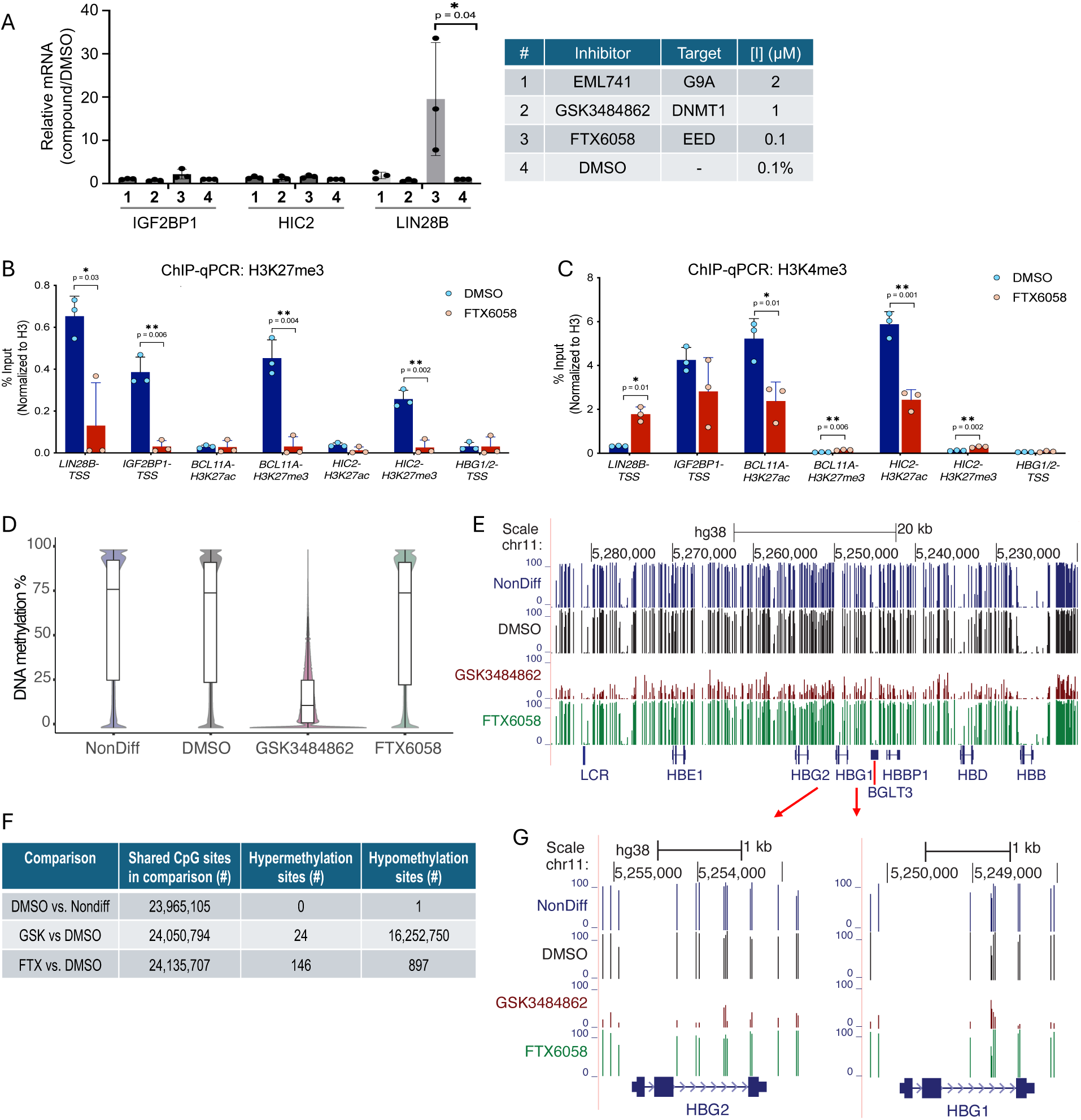
FTX6058 activates *LIN28B* in HUDEP-2 cells. **(A)** Relative mRNA expression levels of *IGF2BP1*, *HIC2*, and *LIN28B* after compound treatment. (**B-C**) ChIP-qPCR analysis of H3K27me3 (panel B) and H3K4me3 (panel C) at gene loci (Supplementary Figure S3) after 7-day FTX treatment (stage 2 cells) normalized to H3. Error bars represent SEM from three different experiments compared to DMSO-treated. **(D) Genome-wide DNA demethylation is restricted to GSK3484862 treatment**. Violin plots showing the distribution of DNA methylation levels in HUDEP-2 cells that are not differentiated (stage 1; NonDiff) and differentiated cells at stage 3 treated with DMSO, GSK, and FTX. Boxplots within the violin plots indicate the first quartile, median, and third quartile methylation values. GSK treatment results in a pronounced global reduction in DNA methylation. (**E**) Genome browser view of the ∼50-kb region on chromosome 11 covering the β-globin locus showing DNA methylation levels (reflected by line heights) at individual CpG sites. (**F**) Number of individual CpG sites that are hypo- or hypermethylated in each condition relative to DMSO. Differentially methylated CpG sites were defined by an absolute methylation difference >25% and a q-value < 0.01. Although far less extensive than with GSK treatment, FTX treatment also induces detectable changes in DNA methylation. (**G**) Genome browser views of the *HBG1* (right) and *HBG2* (left) loci, showing DNA methylation levels (reflected by line heights) at individual CpG sites upstream of, within, and downstream of each gene.

### DNA methylation

To gain insight into genome-wide epigenetic changes induced by these drug treatments, we performed whole genome bisulfite sequencing (WGBS) on stage 1 non-differentiated HUDEP-2 cells and cells at stage 3 after treatment with vehicle (0.1% DMSO), 1 µM GSK, or 0.1 µM FTX. As anticipated, only DNMT1-inhibitor GSK treatment resulted in widespread loss of DNA methylation (Figure 6D; example shown in β-globin locus, Figure 6E). Median DNA methylation levels were 77%, 75%, 12%, and 74% in non-differentiated and stage 3 cells treated with DMSO, GSK, and FTX, respectively.

Consistent with these global trends, differential methylation analysis (absolute difference >25%, q-value < 0.01) revealed minimal DNA methylation changes between non-differentiated and DMSO-treated cells, with only a single hypomethylated CpG site meeting the criteria and no hypermethylated sites detected. In contrast, comparison of GSK-treated versus DMSO-treated cells identified over 16 million hypomethylated CpG sites (Figure 6F), underscoring both the efficacy of the inhibitor and the robust responsiveness of HUDEP-2 cells. Notably, the CpG sites that retained DNA methylation despite GSK treatment were enriched on the X-chromosome (Supplementary Figure S4A). This observation is consistent with prior findings in colorectal cancer cells, in which DNA methylation on the X-chromosome was preferentially retained following prolonged and increased doses of the GSK treatment over several months ^53^.

At the gene level, both *HBG1* and *HBG2* loci are markedly demethylated following GSK treatment (Figure 6G), consistent with the robust reactivation of these genes^54^. Notably, the promoter regions of *HBG1* and *HBG2* do not contain canonical CpG islands, indicating that promoter demethylation is not necessarily required for their re-expression. Indeed, DNA methylation levels were largely unchanged in FTX-treated cells compared with DMSO controls, yet *HBG1* and *HBG2* were strongly re-expressed under FTX treatment. Unexpectedly, FTX treatment was nevertheless associated with localized DNA methylation changes, with approximately 900 CpG sites showing loss of methylation and nearly 150 CpG exhibiting increased methylation. (Supplementary Figure S4B).

### Synergistic effect of three epigenetic inhibitors

We next examined whether the three inhibitors increase fetal globin expression at the cellular level. HUDEP-2 cells were treated with 2 μM EML (G9a-inhibitor), 1 μM GSK (DNMT1-inhibitor), or 0.1 μM FTX (EED-inhibitor) for a total of 9 days and harvested at stage 3 (Figure 5A). Using flow cytometric analysis, we assessed the progression of erythroid differentiation by determining the total proportion of cells positive for erythroid surface markers CD71 and CD235^32^, which showed no notable differences between inhibitor-treated and DMSO-treated cells (Supplementary Figure S2B-C). Interestingly, FTX treatment yielded a higher percentage of cells expressing the early differentiation marker CD105 (∼60%), followed by GSK treatment (∼10%), compared with EML (∼1%) or DMSO treatment (∼1%) (Supplementary Figure S2D). A previous study demonstrated that EED depletion in a murine model of fetal liver cells significantly impeded erythroid maturation^55^. Despite this delayed differentiation, FTX treatment still resulted in an increased proportion of HbF+ cells (∼50%), comparable to GSK (∼60%) and EML (∼40%), all markedly elevated compared with DMSO (∼10%) (Supplementary Figure S2E), suggesting that its mechanism elevates γ-globin independently of normal erythroid maturation.

Next, we evaluated the combined effect of inhibiting the three known epigenetic silencing marks on fetal hemoglobin expression. HUDEP-2 cells were treated with each inhibitor individually or in combination, administered using 10X lower concentrations (0.2 μM EML, 0.1 μM GSK, and 0.01 μM FTX) and harvested at stage 3. The combination of all three inhibitors, even at the 10X lower concentrations, markedly increased the proportion of fetal hemoglobin-expressing cells (∼60%) compared with any single treatment or DMSO (Figure 7A). Cell viability was unaffected by either single or combination treatments (Figure 7B). In addition to expanding the overall HbF+ cell population, the triple-inhibitor combination produced a higher proportion of HbF+ cells with elevated mean fluorescence intensity, suggesting a synergistic effect among the inhibitors. Interestingly, while single treatment with EML (0.2 μM) alone had little effect on the HbF+ cell population (∼15%) (Figure 7A), its inclusion in the triple combination increased the fraction of HbF+ cells with high mean fluorescence intensity (Figure 7C), compared with treatment lacking EML (Figure 7D). These results suggest that G9a inhibition (by EML) synergizes with DNMT1 inhibition (by GSK) and/or EED inhibition (by FTX) in promoting HbF expression. Consistent with the flow cytometry data, the combined treatment with all three compounds at 10X lower doses induced fetal γ-globin protein expression, whereas single treatments did not produce detectable γ-globin by western blots (Figure 7E). Importantly, even at 10 nM, FTX reduced the H3K27me3 mark at stage 3 after 9 days of treatment (Figure 7E).

**Figure 7.**
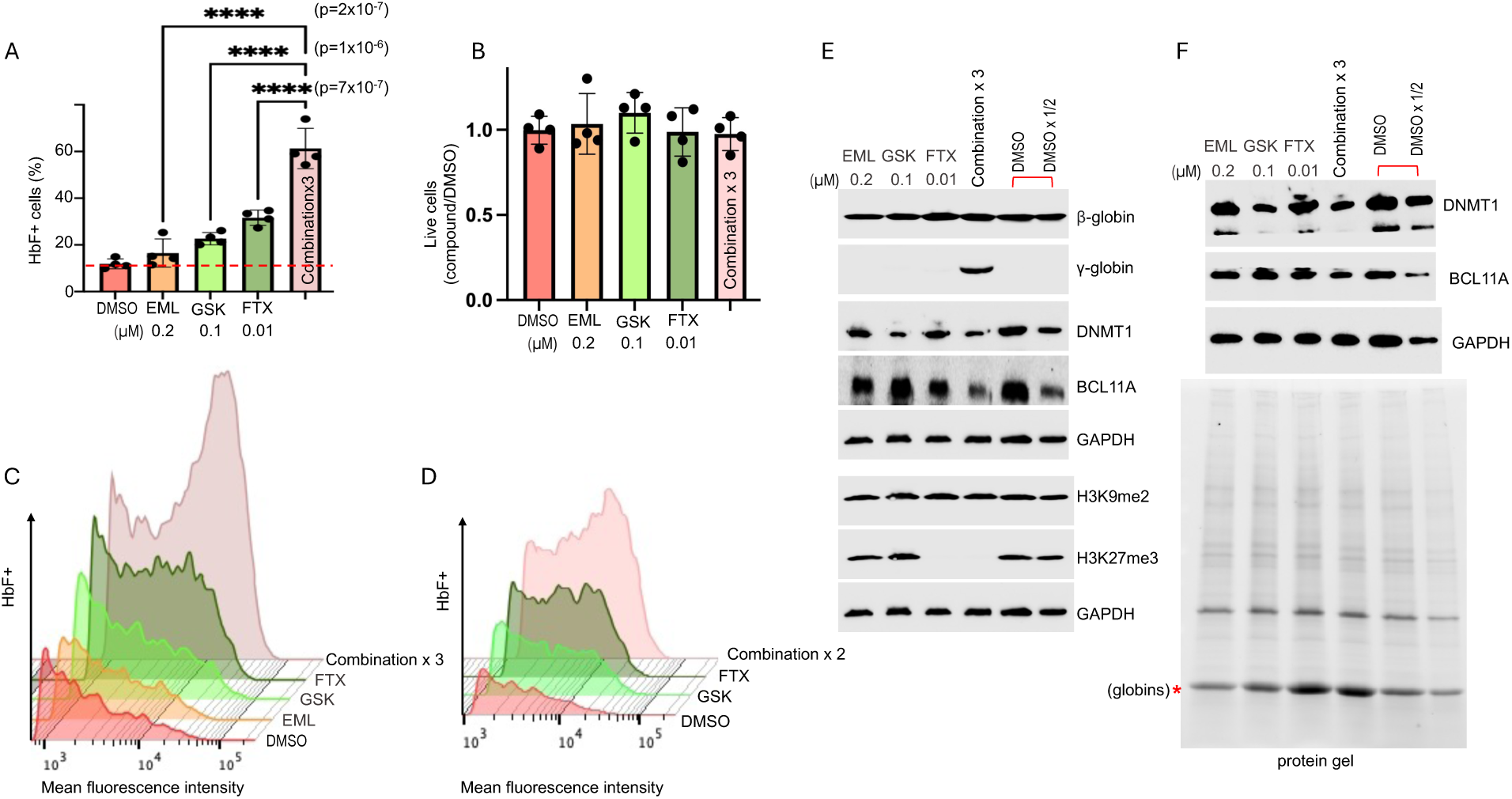
The combined inhibition of three epigenetic silencing marks increases the fetal globin expression in HUDEP-2 cells. HUDEP-2 cells treated with lower doses of individual treatment (0.2 μM EML, 0.1 μM GSK, and 0.01 μM FTX) and together for a total of 9 days treatment (see Figure 5A). (**A**) The percentage of fetal hemoglobin positive (HbF+) cells was determined by flow cytometry. Data represents mean ± SD from N=4 independent experiments. “Combination x3” indicates where all three inhibitors were used together. (**B**) Percentage of live cells quantified using a Bio-Rad cell counter and normalized to the DMSO control. Data represent mean ± SD from N=4 independent experiments. (**C, D**) Representative flow cytometry histogram plot showing HbF+ mean fluorescence intensity (MFI) (X-axis) vs. HbF+ cell count (Y-axis) with and without EML in HUDEP-2 cells, analyzed using FlowJo software. (**E**) Western blot analysis for indicated antibodies and (**F**) stain-free SDS-PAGE protein gel after cells were harvested at stage 3 for a total of 9 days of treatment. The two DMSO lanes as indicated by a bracket are the 2x dilution (1/2) of cell lysates.

### Screen of small molecule inducers of HbF expression

Because genetic ablation of IKZF1 does not fully recapitulate the HbF induction seen with pomalidomide treatment in SCD-derived CD34^+^ HSPCs^28^, we asked whether additional or alternative targets might underlie this effect. To identify small molecules that induce HbF expression in HUDEP-2 cells, we screened 213 pomalidomide- and lenalidomide-derived compounds and quantified HbF+ cells by flow cytometry (Supplementary Table S4).

For the screen, HUDEP-2 cells were treated with 1 μM of compound or DMSO at stages 1 and 2 and harvested at stage 3 (Figure 5A), for a total of 6 days of treatment. Under these conditions, 1 μM pomalidomide increased the percentage of HbF+ cells ∼4.2-fold on average (45%) compared to the DMSO control (11%, p < 0.0001) (Figure 8A). When normalized to internal pomalidomide batch controls, 19 of the 213 compounds (8.8%) induced ≥0.5-fold HbF expression (Figure 8B). These top hits were primarily pomalidomide derivatives with side chain extensions off the 4-amino group, along with a few lenalidomide derivatives (Supplementary Table S4).

**Figure 8.**
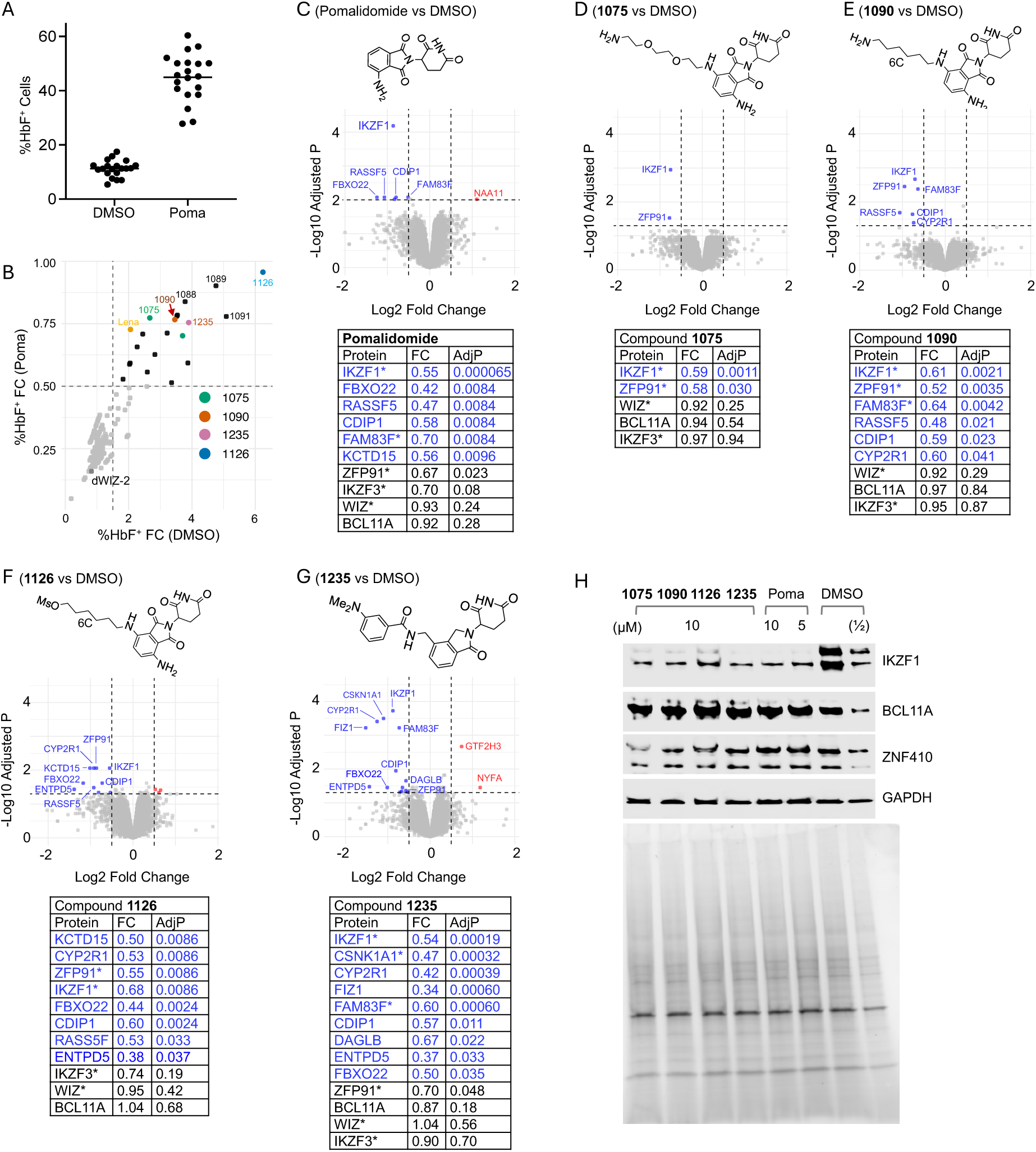
Summary of pomalidomide- and lenalidomide-derivative compound screen for HbF inducers in HUDEP-2 cells and unbiased global proteomic follow-up. For screening, cells were treated at stages 1 and 2 and collected at stage 3 (see Figure 5A). (**A**) Pomalidomide (Poma) treatment (1 µM, 6 days) in HUDEP-2 cells induces a 4-fold increase of %HbF^+^ cells as quantified by flow cytometry (p < 0.0001, two-sided t-test). (**B**) A total of 19 compounds induced HbF levels to at least 50% of the internal pomalidomide positive control, as quantified by flow cytometry. Compounds discussed in text are highlighted in color. Compounds **1088** (4-carbon linker), **1089** (5-carbon linker) and **1091** (7-carbon linker) share chemical structure similarity with **1090**, which contains a 6-carbon linker (Supplementary Table S4). (**C**) Direct targets of pomalidomide in HUDEP-2 cells (indicated by asterisks) include known targets IKZF1 and FAM83F and novel targets FBXO22, RASSF5, CDIP1, and KCTD15 (Supplementary Table S6). (**D**) Compound **1075** led to specific degradation of IKZF1 and ZFP91. (**E**) Compound **1090** led to degradation of a subset of major pomalidomide targets, with increased specificity for ZFP91 and CYP2R1. (**F**) Compound **1126** led to degradation of a subset of pomalidomide targets with increased specificity for CYP2R1 and ZPF91 and decreased specificity for IKZF1. (**G**) Compound **1235**, a derivative of lenalidomide, led to degradation of the known lenalidomide targets (indicated by asterisks) IKZF1, CSNK1A1, and FAM83F, along with the novel targets CYP2R1, FIZ1, CDIP1, DAGLB, and ENTPD5. Two proteins (in red labels) were increased: GTF2H3 (FC = 1.67, adjusted p = 0.0022) and NFYA (FC = 2.27, adjusted p = 0.035). NFYA is necessary for hematopoietic stem cell proliferation^99^. (**H**) Western blot of HUDEP-2 cells at stage 2 treated for 6 hours confirms degradation of IKZF1, shows no detectable degradation of BCL11A, and verifies the presence of ZNF410, which was not detected in the mass spectrometry (Supplementary Table S5). FC = fold change. AdjP = adjusted p-value.

For follow-up protein mass spectrometry, we selected three pomalidomide derivatives: compound **1075**, bearing an amino-terminated six-carbon poly-ethylene-glycol (PEG) chain (pomalidomide-PEG₂-NH₂); compound **1090**, bearing an amino-terminal six-carbon hydrocarbon chain (pomalidomide-C₆-NH₂); and compound **1126**, bearing a mesylate-terminated six-carbon hydrocarbon chain (pomalidomide-C₆-OMs) (Figure 8D-F). We also selected the best-performing hit among lenalidomide derivatives, compound **1235** (Figure 8G).

We used tandem mass tag mass spectrometry^56^ to quantify protein abundance and identify the proteins whose levels are decreased by each compound. HUDEP-2 cells at stage 2 were treated for 6 h with 10 μM compound (N = 2) or 0.1% DMSO (N = 3), and pomalidomide (N = 3) was included as a positive control. As expected, the most significantly decreased protein following pomalidomide treatment was IKZF1 (FC = 0.55, adjusted p = 6.5 x 10^-5^) (Figure 8C and Supplementary Table S5). Previously reported pomalidomide targets in non-erythroid cell types^57^ were also significantly decreased, including FAM83F (Family with Sequence Similarity 83 Member F; FC = 0.70, adjusted p = 0.008) and ZFP91 (Zinc Finger Protein, Atypical E3 Ubiquitin Ligase; FC = 0.67, adjusted p = 0.023) (Figure 8C). Interestingly, several additional proteins not previously described as pomalidomide targets were significantly reduced with adjusted p = 0.0084: FBXO22 (F-Box Protein 22; FC = 0.42), CDIP1 (Cell Death Inducing P53 Target 1; FC = 0.58), and RASSF5 (Ras Association Domain Family Member 5; FC = 0.47). Their potential roles in erythroid function and hemoglobin expression remain unknown, though there is a report of RASSF5 involvement in erythroid differentiation in K562 cells^58^.

Among the three pomalidomide-based compounds (**1075**, **1090**, or **1126**), compound **1075** was tightly selective for IKZF1 and ZFP91 (Figure 8D), while reduced levels of ZFP91 were consistently observed with pomalidomide and all three derivatives (Figure 8C-F). Notably, IKZF1 was identified in a CRISPR-Cas9 screens of ∼1500 DNA binding proteins in HUDEP-2 cells as a regulator of HbF gene expression^9^. Compound **1126,** which induced the highest HbF levels of the screen compounds, most closely phenocopied pomalidomide, with an additional gain of selectivity for CYP2R1 (Figure 8F). ZFP91, identified initially in acute myelogenous leukemia^59^, co-regulates cell survival with IKZF1 in immunomodulatory-resistant T-cell lymphomas^60^.

Compound **1235**, a lenalidomide derivative, led to lower levels of known lenalidomide targets including IKZF1, CSNK1A1 (Casein Kinase 1 Alpha 1), and FAM83F (Figure 8G). Notably FIZ1 (FLT3 Interacting Zinc Finger 1), which is not known to be affected by lenalidomide, showed the most strongly reduced levels (FC = 0.34, p = 6.0 x 10^-4^). While FIZ1 has not been studied in the context of γ-globin repression, its interaction partner FLT3 (FMS Related Receptor Tyrosine Kinase 3) is a regulator of hematopoiesis^61^, and single nucleotide variants (SNPs) in the FLT3 promoter have been associated with increased HbF levels in African sickle cell disease populations^62^.

Using western blot, we validated the degradation of IKZF1 by pomalidomide and the screened compounds using lysates from HUDEP-2 cells at stage 2 treated for 6 hours with the indicated doses of compound (Figure 8H). BCL11A levels were not changed, as seen in the mass spectrometry results. Additionally, we confirmed the expression of the protein ZNF410, which was not detected in our mass spectrometry (Figure 8H).

## DISCUSSION

### BCL11A DNA binding properties

Here, we used three approaches to investigate BCL11A functions: its DNA binding properties, expression dynamics, and its potential for degradation in HUDEP-2 cells. Building on our recent publication on DNA binding by BCL11A^33^, we showed that the two highly similar ZF arrays, ZF2-3 and ZF4-6, recognize all four variants of the TGNCCA motif. These two arrays share 73% sequence identity including 7 out of 8 residues that directly contact DNA bases^33^. A key difference lies in ZF2 Phe388 (which makes a specific hydrophobic contact with the thymine methyl group) and ZF4 Asn753 (which can form hydrogen bonds with variable bases). Among the two TG(A/T)CCA variants most enriched in ChIP peaks, ZF4-6 binds both with high affinity in a strand-specific manner: it engages TGTCCA on the top strand and TGACCA on the complementary strand. To our knowledge, this is the first demonstration of a ZF array binding closely related DNA sequences on opposite strands. The opposite-strand recognition explains the observed conservation of guanines at position 2 on the top strand and of guanines at positions 4 and 5 on the complementary strand (Figure 1C).

We also show that ZF2-3 binds DNA in a manner similar to ZF4-5. The ∼300-residue flexible linker between the two ZF arrays in the same BCL11A molecule enables recognition of motifs that are spaced farther apart and/or arranged in opposite orientations, effectively expanding the binding footprint. Together with recently described N-terminal self-interactions^33,42^, different BCL11A isoforms – including those lacking DNA binding ZF arrays – may form dimers, tetramers, or larger polymers (Figure 9A). Such oligomerization would allow BCL11A to engage multiple clustered motifs, substantially strengthening chromatin occupancy and its repressor activity.

**Figure 9.**
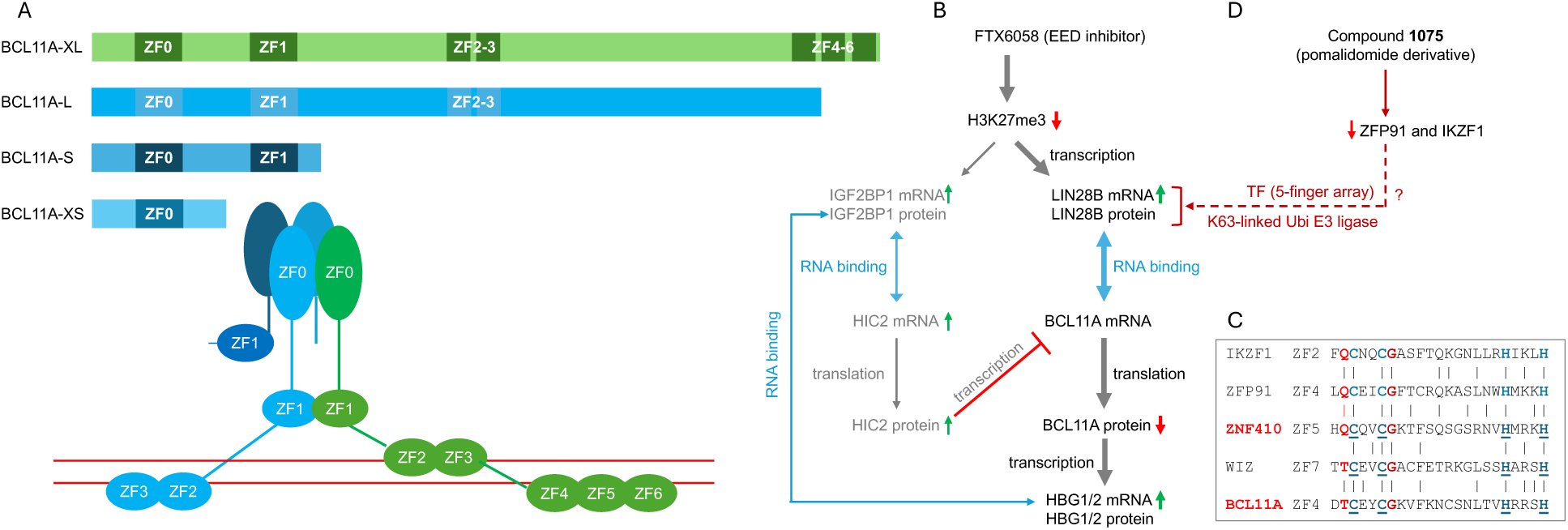
Models of BCL11A regulation across three complementary layers. (**A**) The ZF0-mediated oligomerization assembles different BCL11A splice variants (depicted as different lengths), all of which contain ZF0, enabling BCL11A to form high-order polymers. (**B**) The EED inhibitor FTX6058 reduces BCL11A dosage mainly through LIN28B, which modulates BCL11A translation. (**C**) Sequence alignment of known pomalidomide degrons (IKZF1 and ZFP91) and dWIZ-2 target (WIZ), and the corresponding ZF units of ZNF410 and BCL11A. Zinc-coordinating residues are colored blue, and residues critical for the selectivity of pomalidomide- or lenalidomide-derived degraders are shown in red. (**D**) The pomalidomide-derived compound **1075** induces degradation of ZFP91 and IKZF1, reshaping the regulatory network contributing to HbF induction.

### BCL11A expression dynamics

We used small molecule inhibitors targeting three epigenetic silencing marks. Among them, the EED inhibitor FTX6058 showed the most striking effect, nearly eliminating BCL11A protein levels at 0.1 μM and reducing *BCL11A* mRNA level by half. We reason that this reduction is an indirect consequence of losing H3K27me3 marks. H3K27me3 is normally repressive^63^, so loss of these marks is expected to activate gene transcription rather than repress it. Consistent with this, transcript expression of the RNA binding protein *LIN28B* is increased upon FTX6058 treatment (Figure 9B). LIN28B binds *BCL11A* mRNA and suppresses its translation^48,49^, ultimately reducing BCL11A protein, thereby contributing to HbF induction (Figure 9B).

### Targeted degradation

The final approach we explored was the use of small molecules to glue C2H2 ZF proteins to the cereblon E3 ubiquitin ligase complex, thereby promoting targeted protein degradation^64^. Our initial motivation was to identify additional ZF transcription factors that might be susceptible to pomalidomide- or lenalidomide-derived degraders. We noted that the known cereblon-targeted ZFs do not share a strict consensus amino acid sequence but tend to contain a glycine (G) immediately following the second zinc-ligand cysteine, and often a glutamine (Q) preceding the first zinc-ligand cysteine^65^. ZF5 of ZNF410 contains hallmark residues – Q before the first Cys and G after the second Cys (Figure 9C). A recently reported lenalidomide derivative, dWIZ-2, acts as a molecular glue degrader of WIZ (Widely Interspaced ZF-containing protein)^66^. In this case, ZF7 of WIZ serves as the Cereblon-engaging degron in the presence of dWIZ-2^66^ and exhibits notable sequence similarity to ZF4 of BCL11A – specifically, a threonine (T) preceding the first Cys and a G following the second Cys (Figure 9C). Unexpectedly, ZNF410 was not detected in our mass spectrometry dataset (Supplementary Table S5), and the commercially sourced dWIZ-2 neither depleted WIZ nor induced HbF expression in HUDEP-2 cells (Supplementary Figure S5). However, our compound screen revealed additional insights. Compound **1075** was highly selective for degrading IKZF1 and ZFP91 (Figure 8D), both of which have the characteristic Q-C-xx-C-G motif (Figure 9C).

Although genetic ablation of IKZF1 does not reproduce the HbF induction seen with pomalidomide in SCD-derived CD34^+^ HSPCs^28^, ZFP91 may play a compensatory regulatory role. In immunomodulatory-resistant T-cell lymphomas, ZFP91 supports resistance to IKZF1 loss by regulating genes in multiple pathways such as MAP3K11 (MAP kinase pathway), FZD2 (Wnt signaling pathway), HDAC5 (histone deacetylation), IKBKE (NF-kB pathway), and – most relevant to our discussion – LIN28B^60^. By analogy to the effects of FTX6058, we propose that pomalidomide-derived compounds reduce ZFP91 protein levels in HUDEP-2 cells, leading to altered LIN28B expression/protein levels and ultimately contributing to HbF induction (Figure 9D). ZFP91 might regulate LIN28B through two mechanisms: (1) as a transcription factor via its DNA binding ZF array, influencing mRNA level; and (2) through its K63-linked ubiquitin E3 ligase activity, which ubiquitinates MAP3K14/NIK in HEK293 cells^67^ and could similarly affect LIN28B protein stability.

In summary, our integrated structural, genomic, and chemical-biology approaches reveal three complementary layers of BCL11A regulation. At the DNA level, BCL11A binds short motifs with unexpected strand-specific versatility, enabled by two homologous ZF arrays and a flexible inter-domain linker that together broaden its binding footprint and promote cooperative oligomerization on clustered sites. At the epigenetic level, perturbation of H3K27me3 indirectly reduces BCL11A expression through the activity of RNA-binding regulators such as LIN28B, providing a mechanistic link between chromatin state, RNA biology, and BCL11A dosage. At the protein level, selective degradation of ZF factors such as IKZF1 and ZFP91 by pomalidomide-derived compounds suggests an additional, druggable axis that reshapes BCL11A’s regulatory network. Collectively, these findings define multiple mechanistic entry points—structural, transcriptional, and post-translational—that can be exploited to modulate BCL11A activity for therapeutic HbF reactivation.

## Methods

Expression and purification of BCL11A ZF2-3 and ZF4-6 fragments proceeded as previously discussed ^33^. Oligonucleotides were purchased from Integrated DNA Technologies (IDT). Chemical compounds GSK3484862 (HY-135146), FTX6058 (HY-139400), pomalidomide (HY-10984) and lenalidomide (HY-A0003) were purchased from MedChemExpress. Compound EML741 was synthesized and prepared as described^17^. Compounds were dissolved in 100% dimethyl sulfoxide (DMSO), aliquoted, and stored at −80°C prior to use.

### DNA binding assays

We used two biophysical methods for protein-DNA binding assays using FAM-labeled DNA duplex: (1) fluorescence polarization (FP)-based DNA binding assay and (2) electrophoretic mobility shift assay (EMSA).

A total volume of 40 µL was used for each reaction, containing 20 nM FAM-labeled DNA duplex and different concentrations of BCL11A protein. The reactions were carried out in buffer with 20 mM Tris-HCl (pH 7.5), 150 mM NaCl, 5% glycerol, and 0.5 mM TCEP, followed by incubation on ice for 20 min. Fluorescence polarization measurements were taken using a Synergy Neo2 Multi-Mode Reader (BioTek). Data were processed with GraphPad Prism 10, and the dissociation constant (K_D_) was obtained by fitting to the equation: [mP]= [maximum mP] × [C] / (K_D_ + [C]) + [baseline mP], where mP denotes the observed milli-polarization and [C] represents the protein concentration.

After the fluorescence polarization measurements, 10 µL from each reaction mixture was analyzed by electrophoresis on an 8% native polyacrylamide gel. The gel was run at a constant 150 V for 25 min in ice-cold 0.5× TBE buffer, followed by imaging with a Typhoon 9410 scanner (GE Healthcare).

### BCL11A binding motif analysis

ChIP-seq datasets (GSM2771529 and GSM2771530)^30^ were downloaded from Cistrome (v3.0)^68^. CUT&RUN datasets^31^ were obtained from GEO (GSE104676; NarrowPeak files GSM2805366, GSM2805367, GSM2805368). Peak sequences were extracted using the Fasta and SeqIO modules from Pyfaidx and Biopython, respectively. Human genome hg38 was used for ChIP-seq and hg19 for CUT&RUN, matching the original analyses. TGNCCA sequences and their reverse complements were counted in each replicate to obtain observed motif frequencies. Expected motif counts were derived using first-order Markov models of nucleotide composition generated from the peak sequences of each replicate using MEME Suite (v5.5.5)^69,70^, providing the probability of each base occurring given the preceding base. Transition probabilities were used to estimate the expected frequency of each motif (and its reverse complement) within each peak, and expected counts were summed across peaks. Global enrichment was assessed using a two-sided exact Poisson test in R. Peak annotation for both ChIP-seq and CUT&RUN data was performed using HOMER (v5.1)^41^ with the appropriate reference genomes.

### Crystallography

Oligonucleotides were annealed in a buffer containing 50 mM NaCl and 10 mM Tris-HCl pH 7.5. BCL11A proteins were concentrated to 4-6 mM and the annealed DNA duplex was added at a 1.25:1 molar ratio of protein to DNA, yielding a final complex concentration of ∼1.6 mM. The mixture was incubated overnight at 4 °C. Crystallization trials were performed using commercially available screening kits and an Art Robbins Gryphon Crystallization Robot to set up sitting-drop vapor diffusion experiments (0.2 μL protein-DNA complex + 0.2 μL reservoir solution).

Crystallization plates containing several hundred screening conditions were incubated at 19°C. The conditions that yielded crystals, from which X-ray diffraction were collected, were summarized in Supplementary Table S7. Crystals of BCL11 ZF4-6 with TGTCCA-containing oligonucleotides grew under many conditions within 1-5 days, whereas crystals of BCL11A ZF4-6 with the TGGCCA oligonucleotide formed under fewer conditions.

Diffraction-quality crystals of BCL11A ZF2-3 bound to DNA appeared under far fewer conditions, with some producing showers of very small crystals at 19° C. Crystal grown with 12-bp TGTCCA-containing oligonucleotide yielded a form belonging to space group *P*2_1_2_1_2. For conditions that produced microcrystals, additional sitting-drop experiments were setup and incubated at 4 °C. Several weeks later, large crystals of a distinct morphology were observed at 4°C using the same oligonucleotide, yielded crystals in space group *C*222_1_.

All crystals were harvested using a nylon or Litho loop and flash-frozen in liquid nitrogen after momentary soaking in reservoir solution supplemented with 20% (v/v) ethylene glycol when the original condition lacked a cryoprotectant. All birefringent crystals were shipped to beamline 17-ID-1 (AMX) at the National Synchrotron Light Source II (NSLS-II)^71^, Brookhaven National Laboratory, and maintained in a 100 K cryostream for screening and, when adequate diffraction was observed, subsequent data collection. Diffraction data were processed with the autoPROC toolbox^72^, which employs XDS^73^ for data reduction. Intensities from images representing 360° of crystal rotation were merged using POINTLESS^74^ and AIMLESS^75^. However, the Xtriage module^76^ within the PHENIX package indicates that all datasets from crystals containing BCL11A ZF4-6 and both TGTCCA oligonucleotides (12-bp and 20-bp) exhibited severe anisotropy, which hindered molecular replacement and prevented the generation of interpretable electron density maps. These datasets were subsequently reprocessed using STARANISO software (Cambridge, United Kingdom: Global Phasing Limited) (http://staraniso.globalphasing.org/cgi-bin/staraniso.cgi), which performs anisotropic cutoffs of merged intensity data, applies Bayesian estimation of structure amplitudes, and corrects for anisotropy. The STARANISO-processed datasets were then used for molecular replacement, map generation, and refinement.

The structures of the BCL11A ZF4-6 and DNA complexes were determined by molecular replacement using the PHASER module^77^ in the PHENIX suite, with PDB 9E6R serving as the initial search model. For the *P*2_1_2_1_2 dataset of BCL11A ZF2-3 with DNA, an Alphafold3 model of BCL11A ZF2-3 was used and the resulting solution was subsequently used as the search model for the *C*222_1_ form. All structures were refined using phenix.refine^78,79^. A randomly selected 5% subset of reflections was excluded from refinement and used to monitor R-free values^80^ for cross-validation. Manual model building and corrections were carried out in COOT^81,82^ between refinement cycles. After each round of refinement, structure quality was carefully evaluated, and final validation was performed using the PDB validation server^83^. Molecular graphics were generated using PyMol (Schrödinger, LLC).

### HUDEP-2 cell culture and erythroid differentiation

HUDEP-2 cells were cultured as previously described^14^. Cells were maintained in expansion medium (StemSpan^TM^ SFEM, STEMCELL #09650) supplemented with 50 ng/mL stem cell factor (SCF) (STEMCELL #78062), 1 μg/mL doxycycline (Sigma-Aldrich #D9891), 0.4 μg/mL dexamethasone (Sigma-Aldrich #D4902), 3 IU/mL erythropoietin (STEMCELL #78007), 2% penicillin–streptomycin (Gibco #15140148), and 1% L-glutamine (STEMCELL #07100). For stage 1, cells were expanded for 3 days.

Erythroid differentiation followed a two-phase protocol^15^. Erythroid differentiation medium (EDM) consisted of Iscove’s Modified Dulbecco’s Medium (IMDM) (Sigma-Aldrich #51471C) supplemented with 330 μg/mL holo-transferrin (Sigma-Aldrich #T0665), 10 μg/mL insulin (Sigma-Aldrich #I9278), 2 IU/mL heparin (Sigma-Aldrich #H3149), 5% fetal bovine serum (FBS) (Sigma-Aldrich #F4135), 3 IU/mL erythropoietin, 2% penicillin–streptomycin and 1% L-glutamine. For differentiation-growth phase (4 days), EDM was further supplemented with 100 ng/mL SCF and 1 μg/mL doxycycline. For differentiation-maturation phase (2 days), EDM contained only 1 μg/mL doxycycline. All cultures cells were maintained at 37°C with 5% CO_2_. Compounds in 0.1% DMSO (final) were added in fresh medium at day 0 (stage 0), day 3 (stage 1), and day 7 (stage 2) (Figure 5A).

### Flow cytometry

HbF expression was assessed by intracellular staining. Approximately 500,000 HUDEP-2 cells in stage 3 were harvested, fixed and permeabilized using the Cyto-Fast^TM^ Fix/Perm Buffer set (BioLegend #426803) for 20 min at room temperature in the dark, following the manufacturer protocol. Cells were incubated with FITC-conjugated anti-human fetal hemoglobin antibody (REAfinity^TM^, clone REA533; Miltenyi Biotec #130-120-065) for 20 min at room temperature in the dark.

To assess erythroid differentiation, cells were stained with PE/Cyanine7-conjugated anti-CD71 (BioLegend #334112), APC-conjugated anti-CD235a (BioLegend #349114), and PE-conjugated anti-CD105 (BioLegend #323205). Corresponding isotype controls antibodies used were PE/Cyanine7 Mouse IgG2a (BioLegend #400231), APC Mouse IgG2a (BioLegend #400221), and PE Mouse IgG1 (BioLegend #400113). Flow cytometry data were acquired on a Gallios Flow Cytometer (Beckman Coulter) and analyzed using FlowJo v10.10.0 (Becton Dickson).

### Western blots

Cells were harvested, washed twice with 1x PBS, lysed in 3x SDS (sodium dodecyl sulfate) cracking buffer, and heated at 95°C for 5 min. Whole cell lysate were resolved on 4–20% Mini-Protean TGX stain-free gel (Bio-Rad #4568096) and transferred to Immun-Blot^R^ Low PVDF Fluorescence membranes (Bio-Rad #1704274) using the Trans-Blot Turbo Transfer System (Bio-Rad). Membranes were blocked in 5% nonfat dry milk (Bio-Rad #1706404) in TBST for 1 h at room temperature. The comparison SDS gels were BioRad stain-free imaging taken before transferring to blot on ChemiDoc Go Imaging System (Bio-Rad).

Membranes were incubated with primary antibodies overnight at 4°C. Primary antibodies included DNMT1 (Cell Signaling Technology - CST #5032), H3K9me2 (CST #4658), H3K27me3 (CST #9733s), H3 (CST #14269), EED (CST #51673), GAPDH (CST #5174), β-globin (37-8) (Santa Cruz #21757), γ-globin (51-7) (Santa Cruz #21756), BCL11A (Novus Biologicals #NB600-261), ZNF410 (Proteintech #14529-1-AP), IKZF1 (CST #14859) and actin (Sigma-Aldrich #A2228). Secondary antibodies were HRP-conjugated anti-rabbit-IgG (CST #7074) and HRP-conjugated anti-mouse-IgG (ImmumoReagents GtxMu-003-FHRPX) were applied for 1 h at room temperature. Signals were detected using Clarity^TM^ Western ECL (Bio-Rad #1705061) on a ChemiDoc Go Imaging System (Bio-Rad).

### RNA isolation and RT-qPCR

Cells at stage 2 differentiation were harvested and pelleted, and total RNAs were extracted using TRIzol (Invitrogen #15596-018) following the manufacturer’s protocol. Genomic DNA was removed, and cDNA sunthesized using the SuperScript IV VILO Master Mix with ezDNase (Invitrogen #11766050). RT-qPCR was performed in technical duplicates using PowerTrack SYBR Green Master Mix (Applied Biosystems #A46109) with 10 ng of cDNA per 20 uL reaction. Primers sequences are listed in Supplementary Table S8. The mRNA levels normalized to Importin 8 (IP08), and fold changes were calculated using the 2(-ΔΔCt) method^84^.

For statistical testing, raw Ct values were converted to ΔCt (Ct_target − Ct_reference). ΔCt values were analyzed using a one-way mixed-effects model (repeated measures by experiment) with treatment as the fixed effect and experiment as a random effect; comparisons to the DMSO control were performed using Dunnett’s correction. Fold changes reported in Figure 5I–5J are 2^−ΔΔCt relative to DMSO.

### ChIP–qPCR analysis of histone modifications

Chromatin immunoprecipitation (ChIP) assays were performed at the MD Anderson Cancer Center Epigenomics Profiling Core as described previously^85,86^ with some modifications. Briefly, HUDEP-2 cells were treated with DMSO or 0.1 μM FTX6058 for 7 days (stage 2 cells) and crosslinked in 1% formaldehyde for 8 min, followed by quenching with 125 mM glycine for 5 min at room temperature. Nuclei were isolated and lysed to prepare chromatin, which was then subjected to sonication using a Bioruptor Pico (Diagenode) to generate DNA fragments ranging 200-600 bp. Lysates were cleared by centrifuged at 16,000g for 10 min at 4°C.

The resulting supernatants were incubated overnight at 4°C with antibodies against H3K4me3 (Abcam #ab213224), H3 (Abcam #ab1791), or H3K27me3 (Diagenode #C15410195), pre-conjugated to Dynabeads Protein A beads (Invitrogen #10001D). Next day, immunocomplexes were collected using a DynaMag-2 magnet (Invitrogen #12321D), washed extensively according to the manufacturer’s protocol, and reverse crosslinked, followed by DNA extraction. Enrichment of target genomic regions were quantified by SYBR Green-based real-time quantitative PCR (qPCR) using gene-specific primers (Supplementary Figure S3), which were designed based on recent published CUT&RUN and CUT&Tag datasets available in GEO (GSE292698)^87^. Histone-mark enrichment at each gene locus was calculated as percent input and normalized to total H3.

### Whole Genome Bisulfite Sequencing (WGBS)

Four samples were prepared for WGBS analysis. Stage 1 cells and DMSO-treated, stage 3 cells were collected in bulk without sorting. For GSK3484862 (1 μM) or FTX6058 (0.1 μM) treated HUDEP-2 cells, stage 3 cells were harvested, fixed and permeabilized using the Cyto-Fast^TM^ Fix/Perm Buffer Set (BioLegend #426803) for 20 min at room temperature in the dark, followed by staining with FITC-conjugated anti-human fetal hemoglobin antibody (REAfinity^TM^, clone REA533; Miltenyi Biotec #130-120-065) for 20 min. HbF+ cells were isolated using a BD FACSAria SORP sorter.

Proteins were digested from sorted HbF+ cells by overnight incubation at 50 °C in digestion buffer containing proteinase K (Invitrogen #K182001). Genomic DNA was extracted using the PureLink Genomic DNA Mini Kit (Invitrogen #K182001). Between 10–100 ng of genomic DNA was spiked with 0.5% lambda DNA and subjected to bisulfite conversion using the EZ DNA Methylation-Lightning Kit (Zymo Research #5031). Sequencing libraries were prepared from bisulfite-converted DNA using the Accel-NGS Methyl-Seq DNA Library Kit (Swift Biosciences #30024). Final libraries were pooled and sequenced on an Illumina NovaSeq X plus 25B platform (2 x 150 bp) in paired-end 150/8/8/150 configuration.

Adapter sequences and low-complexity reads were removed using Trim Galore! (v0.6.7) (https://www.bioinformatics.babraham.ac.uk/projects/trim_galore/), cutadapt (v4.1)^88^, and an in-house bash script. Filtered reads were aligned to the human reference genome (hg38) using the bisulfite converted read mapper Bismark (v0.16.3)^89^ and Bowtie2 (v2.1.0)^90^. Approximately 79-80% of read pairs uniquely mapped to the human genome. After deduplication with the Bismark’s deduplicate_bismark script, 469-529 million uniquely mapped read pairs were retained per sample, yielding an average CpG coverage of 36x-42x.

CpG methylation levels were extracted using the bismark_methylation_extractor script and an in-house Perl script. CpG sites with ≥10x read coverage were considered qualified for further analysis. Each sample contained 24.7–25.0 million qualified CpG sites, covering 84.0-84.9% of the human genome. Differential methylation analysis was performed using the R/Bioconductor package methylKit (v1.34.0)^91^, using only qualified CpG sites present in both samples of each comparison. CpG sites with an absolute methylation difference ≥25% and a q-value ≤0.01 were defined as differentially methylated.

### Chemical synthesis of pomalidomide- and lenalidomide-derivatives

Compounds **1075**, **1090** and **1126** (pomalidomide-derivatives) were prepared according to reported procedures^92,93^. The lenalidomide-derivative **1235** [3-(dimethylamino)-*N*-((2-(2,6-dioxopiperidin-3-yl)-1-oxoisoindolin-4-yl)methyl)benzamide] was synthesized as follows (Supplementary Figure S6A): lenalidomide (5.0 mg, 0.018 mmol) was dissolved in DMSO (1 mL), then 3-(dimethylamino)benzoic acid (10.0 mg, 0.061 mmol), HOAt (8.2 mg, 0.060 mmol), EDCI (11.5 mg, 0.060 mmol), and NMM (40 μL, 0.36 mmol) were added. The mixture was stirred at room temperature for 1 h, then purified by prep-HPLC to give compound **1235** as yellow solid (6.5 mg, 67% yield). ^1^H NMR (400 MHz, DMSO-*d*_6_) δ 11.02 (s, 1H), 8.99 (t, *J* = 5.9 Hz, 1H), 7.63 (d, *J* = 7.3 Hz, 1H), 7.58 – 7.47 (m, 2H), 7.26 (t, *J* = 7.9 Hz, 1H), 7.21 – 7.13 (m, 2H), 6.88 (dd, *J* = 8.1, 2.9 Hz, 1H), 5.15 (dd, *J* = 13.3, 5.1 Hz, 1H), 4.61 – 4.38 (m, 4H), 3.00 – 2.84 (m, 7H), 2.69 – 2.57 (m, 1H), 2.39 (dq, *J* = 13.3, 4.5 Hz, 1H), 2.09 – 1.98 (m, 1H) (Supplementary Figure S6B). ^13^C NMR (101 MHz, DMSO) δ 173.36, 171.52, 168.59, 167.46, 150.71, 140.56, 135.42, 135.25, 132.09, 131.12, 129.34, 128.81, 122.10, 115.62, 115.45, 111.48, 52.07, 46.80, 40.64, 39.98, 31.67, 23.09 (Supplementary Figure S6C). HRMS (ESI-TOF) m/z: [M+H]^+^ calcd for C_23_H_25_N_4_O_4_, 421.1870; found: 421.1878.

All commercial reagents and solvents were used without further purification. Flash column chromatography was performed on a Teledyne ISCO CombiFlash Rf+ instrument with UV detector 220/254/280 nm. Normal-phase chromatography was conducted on silica gel using hexane/ethyl acetate or dichloromethane/methanol as eluent. Reverse-phase chromatography using HP C18 RediSep Rf columns with a gradient from 10% of MeOH in H_2_O containing 0.1% TFA to 100% of MeOH.

All final compounds were purified with preparative high-performance liquid chromatography (HPLC) on an Agilent Prep 1290 infinity II series (UV detection at 220/254/280 nm, 40 mL/min) (for compound **1235**, see Supplementary Figire S6D). Samples were injected onto a Phenomenex Luna 750 x 30 mm, 5 μm C18 column and eluted with a gradient from 10% acetonitrile in H_2_O containing 0.1% TFA to 100% acetonitrile.

The purity of all compounds was assessed using an Agilent 1200 series system with DAD detector and a 2.1 mm x 150 mm Zorbax 300SB-C18 5 μm column for chromatography and high-resolution mass spectra (HRMS). HRMS data were acquired in positive ion mode on either an Agilent G6230BA Accurate Mass TOF or an Agilent G1969A API-TOF with an electrospray ionization (ESI) source. Samples (0.8-2 μL) were injected onto a C18 column at room temperature and eluted at 0.4-0.8 mL/min using water containing 0.1% formic acid (solvent A) and acetonitrile containing 0.1% formic acid (solvent B).

Nuclear magnetic resonance (NMR) spectra were collected on a Bruker DRX instrument (^1^H NMR 400 MHz, ^13^C NMR 101 MHz). Chemical shifts for all compounds are reported in parts per million (ppm, δ) with standard annotations for multiplicity (s = singlet, d = doublet, t = triplet, q = quartet, m = multiplet), coupling constant (J values in Hz), and integration. All final compounds tested were > 95% pure by the HPLC methods described above.

### Molecular glue degrader compound screen

Compound (1 μM) or DMSO (0.1%) were added to HUDEP-2 cells at stages 1 and 2 and harvested for flow cytometry at stage 3 (Figure 5A). Cells were fixed, permeabilized, and stained for intracellular HbF as described above (see Method on Flow cytometry). Flow cytometry data were acquired using LSRFortessa SORP (Becton Dickson), data were analyzed by FACSDiva Software (Becton Dickson), and data were plotted using FlowJo v10.10.0 (Becton Dickson).

### MSPC001383 Proteome Profiling

HUDEP-2 cells were lysed in 8M urea buffer, reduced/alkylated and digested using LysC and Trypsin proteases at 37°C overnight. The peptide cleanup was performed on Bravo Automated Liquid Handling Platform (Agilent) using the C18-5µl tips (Agilent, 5190-6532). The peptides were quantified using the Pierce™ Quantitative Colorimetric Peptide Assay (Thermo Scientific 23275). 20µg peptide per sample were labeled with TMTpro 16-plex isobaric label reagent (Thermo Fisher Scientific, A44521) according to manufacturer’s protocol. The high-pH offline fractionation was carried on Agilent 1260 Infinity II system to generate 24 peptide pools and acidified with final concentration of 0.1% formic acid (FA). The deep-fractionated peptide samples were analyzed on Vanquish Neo UHPLC system (Thermo Fisher Scientific, San Jose, CA) coupled to Orbitrap Eclipse (Thermo Fisher Scientific, San Jose, CA) mass spectrometer. The peptide separation was carried out on 20cm X 75µm I.D. analytical column (Reprosil-Pur Basic C18aq, Dr. Maisch GmbH, Germany) at 200 nl/min flow rate for 110 min gradient time. The MS1 was done in Orbitrap (120,000 resolution, scan range 375-1500 m/z, 50 ms Injection time) followed by MS2 in Orbitrap at 30,000 resolution (HCD 38%) with TurboTMT algorithm. Dynamic exclusion was set to 20 sec and the isolation width was set to 0.7 m/z. The mass spectra were searched using MSFragger (v3.5). The reverse decoys and common contaminants were added to the GENCODE human protein database (v42) using Philosopher^94^. Peptide validation was performed using semi-supervised learning procedure in Percolator^95^ as implemented in MokaPot^96^. The MS raw data processing, quantification and differential analysis was carried out as described before for proteomics profiling^97^. The gene product inference and iBAQ-based quantification was carried out using the gpGrouper algorithm^98^ to calculate peptide peak area (MS1) based expression estimates. The median normalized and log2 transformed iBAQ values were used for data analysis. The differentially expressed proteins were calculated using the moderated t-test to calculate p-values and log2 fold changes in the R package limma. The False Discovery Rate corrected pValue was calculated using the Benjamini-Hochberg procedure.

## DATA AVAILABILITY

The coordinates and structure factor files of the X-ray structures of the BCL11A in complexes with DNA have been deposited to PDB and released under accession numbers 9YLL for ZF4-6 and TGTCCA in 20-bp duplex, 9YLM for ZF4-6 and TGTCCA in 12-bp duplex, 9YLN for ZF4-6 and TGGCCA in 12-bp duplex, 9YLO for ZF2-3 and TGTCCA in *P*2_1_2_1_2 space group, 9YLP for ZF2-3 and TGTCCA in *C*222_1_ space group. The mass spectrometry proteomics data have been deposited to the ProteomeXchange Consortium via the PRIDE partner repository with the dataset identifier PXD072312. WGBS data with accession number GSE316022 can be accessed through secure token: gjivmcokppwlrax.

## ACKNOWLEDGEMENTS

We thank Pamela Whitney for help with flow cytometry, in part using Epigenetics and Molecular Carcinogenesis Flow Cytometry Core Facility and the Flow Cytometry and Cellular Imaging Facility (FCCIF) at the University of Texas MD Anderson Cancer Center (MDACC). The use of FCCIF is partially supported by the National Cancer Institute grant P30CA016672 and a Shared Instrumentation Award from the Cancer Prevention Research Institution of Texas (CPRIT). We thank Antrix Jain, Shirley Wang and Anna Malovannaya (Baylor College of Medicine) for performing mass spectrometry. The use of Baylor College of Medicine Mass Spectrometry Proteomics Core (RRID:SCR_027015) is supported by the Dan L. Duncan Comprehensive Cancer Center NIH award (P30 CA125123) and CPRIT Core Facility Award (RP210227). We thank Xiangpeng Kong (New York University) for assistance of access to 17-ID-1 beamtime and the beamline scientists of the National Synchrotron Light Source (NSLS) II, Brookhaven National Laboratory. The use of NSLS II resources 17-ID-1 was supported by the U.S. Department of Energy under contract DESC0012704. The use of MDACC Epigenomics Profiling Core is partially supported by the institutional funds. M.Y. thanks Cristian Coarfa (Baylor College of Medicine) for bioinformatics guidance.

## AUTHOR CONTRIBUTIONS

M.Y. performed bioinformatics (motif) analysis and compound screens, contributed to manuscript writing. P.D. performed epigenetic inhibitions, flow cytometry, western blots, RNA isolation and RT-qPCR, contributed to manuscript writing. J.R.H. performed crystallographic analyses. J.Z. and J.L. performed FP and EMSA binding assays, and protein purifications. T.H. and Y.H. performed WGBS-bisulfite conversion and library preparation. Y.L. and M.R.E. performed WGBS DNA methylation analysis. P.A.I. and A.K.J. performed ChIP-qPCR and analysis. G.S. provided compound EML741. Y.X. and J.J. provided pomalidomide and lenalidomide derivatives. R.M.B. participated in discussions, and in preparing the manuscript. Y.H. contributed funding acquisition. X.Z. contributed to discussions, supervision and project administration. X.C. designed and organized the study, contributed to manuscript writing and editing, conceptualization, and funding acquisition.

## Footnotes

1 https://www.fda.gov/news-events/press-announcements/fda-approves-first-gene-therapies-treat-patients-sickle-cell-disease

2 https://www.nytimes.com/2024/10/21/health/sickle-cell-disease-gene-therapy-patient.html

3 https://doi.org/10.1182/blood-2020-142056; https://doi.org/10.1182/blood-2020-141145; https://doi.org/10.1182/blood-2023-190115; https://pmc.ncbi.nlm.nih.gov/articles/PMC9429145/; https://doi.org/10.1182/blood-2025-1157

