## supplementary tables and figures for "Regulation of BCL11A DNA binding and expression in human erythrocyte precursor HUDEP-2 cells"

Table S1. Summary of observed and expected TGNCCA motif counts from BCL11A ChIP-seq (Martyn et al. 2018; hg38) and CUT&RUN (Liu et al. 2018; hg19) data and Markov models used to generated expected motif count.

Table S2. List of validated BCL11A target genes from Mehta et al. 2022 and the number of peaks from Martyn et al. 2018 and Liu et al. 2018 associated with each peak.

Table S3. Proportion BCL11A target genes from Mehta et al. 2022 compared to the proportion of total genes across chromosomes.

Table S4. List of 213 compounds with chemical structure, molecular weight, and Fold Change

Table S5. Unique to gene peptide counts for selected proteins as detected by TMT mass spectrometry proteomics in HUDEP-2 cells.

Table S6. List of significantly differentially expressed proteins compared to DMSO (adjP < 0.05) for compounds assessed by TMT mass spectrometry. Additional other relevant proteins (WIZ, BCL11A, etc.) also listed.

The following tables appear in Methods:

Table S7. Crystallographic Table

Table S8. Primers used for qPCR

Table S7.

Summary of X-ray data collection at beamline 17-ID-1, NSLSII ($\lambda$=0.92010Å) and structure refinement statistics

| Protein BCL11A | ZF4-6 | ZF4-6 | ZF4-6 | ZF2-3 | ZF2-3 |
|---|---|---|---|---|---|
| DNA (5'-3') | GAA**TGTCCA**TCTTGCAAAAC | GAA**TGTCCA**TCTC | GAA**TGGCCA**TCTC | GAA**TGTCCA**TCT | GAA**TGTCCA**TCT |
| (3'-5') | CTTACAGGTAGAACGTTTTG | GCTTACAGGTAGA | GCTTACCGGTAGA | CTTACAGGTAGA | CTTACAGGTAGA |
| PDB Code | 9YLL | 9YLM | 9YLN | 9YLO | 9YLP |
| Date Collected | 11/20/2024 | 02/14/2025 | 03/20/2025 | 03/20/2025/ | 05/17/2025 |
| Space group | $P4_32_12$ | $P4_12_12$ | $P4_12_12$ | $P2_12_12$ | $C222_1$ |
| Cell dimensions (Å) | 59.23, 59.23, 247.1 | 56.62, 56.62, 154.8 | 56.97, 56.97, 155.9 | 122.4, 29.96, 39.11 | 36.90, 69.55, 129.2 |
| $\alpha, \beta, \gamma$ (°) | 90, 90, 90 | 90, 90, 90 | 90, 90, 90 | 90, 90, 90 | 90, 90, 90 |
| Resolution (Å) | 34.90-2.38 (2.76-2.38) | 31.95-1.91 (2.15-1.91) | 31.8-2.15 (2.18-2.15) | 29.1-1.97 (2.00-1.97) | 34.78-2.04 (2.08-2.04) |
| [a] $R_{merge}$ | 0.154 (2.21) | 0.093 (2.18) | 0.200 (5.85) | 0.198 (2.74) | 0.093 (1.81) |
| $R_{pim}$ | 0.032 (0.478) | 0.028 (0.622) | 0.043 (1.21) | 0.065 (0.847) | 0.027 (0.522) |
| $CC_{1/2}$ | 0.998 (0.646) | 0.988 (0.554) | 0.977 (0.430) | 0.990 (0.606) | 0.999 (0.355) |
| [b] <I/σI> | 13.7 (1.9) | 14.1 (1.4) | 10.9 (0.9) | 6.7 (1.4) | 11.9 (1.5) |
| Completeness (%) | 46.5 (6.7) [e]<br>91.3 (34.90-3.23) [e] | 71.0 (12.1) [e]<br>99.6 (31.95-2.26) [e] | 100 (100) | 98.4 (100.0) | 100 (100) |
| Redundancy | 24.7 (21.9) | 13.0 (13.0) | 23.4 (24.1) | 10.4 (11.3) | 13.3 (12.9) |
| Observed reflections | 219,053 (9,705) | 189,522 (9,444) | 346,555 (17,657) | 110,452 (5,973) | 145,626 (6,903) |
| Unique reflections | 8,878 (444) | 14,526 (726) | 14,815 (732) | 10,634 (529) | 10,930 (535) |
| **Refinement** | | | | | |
| Resolution (Å) | 2.38 | 1.91 | 2.15 | 1.97 | 2.04 |
| No. reflections | 8,865 | 14,504 | 14,720 | 10,554 | 10,903 |
| [c] $R_{work}$ / [d] $R_{free}$ | 0.197 / 0.248 | 0.204 / 0.243 | 0.202 / 0.247 | 0.215 / 0.242 | 0.208 / 0.243 |
| No. Atoms | | | | | |
| Protein | 668 | 667 | 665 | 427 | 453 |
| DNA | 814 | 527 | 512 | 486 | 486 |
| Zn | 3 | 3 | 3 | 2 | 3 |
| Solvent | 7 | 73 | 47 | 45 | 47 |
| B Factors (Å$^2$) | | | | | |
| Protein | 67.3 | 49.1 | 68.6 | 44.0 | 55.0 |
| DNA | 80.5 | 57.0 | 75.5 | 58.3 | 58.7 |
| Zn | 63.8 | 36.5 | 52.4 | 31.1 | 56.8 |
| Solvent | 29.1 | 49.4 | 63.2 | 46.9 | 59.1 |
| **R.m.s. deviations** | | | | | |
| Bond lengths (Å) | 0.006 | 0.004 | 0.006 | 0.003 | 0.005 |
| Bond angles (°) | 0.7 | 0.6 | 0.8 | 0.5 | 0.7 |
| Crystallization conditions | 20% (w/v) PEG 3350<br>0.2 M NaCl | 28% (w/v) PEG 4000<br>0.1 M sodium citrate<br>(pH 5.2)<br>0.2 M ammonium acetate | 28% (w/v) PEG 4000<br>0.1 M sodium citrate<br>(pH 5.2)<br>0.2 M ammonium acetate | 10% (w/v) PEG 20000<br>20% PEG MME 500<br>0.1M Bicine-TRIS<br>(pH 8.5)<br>60 mM NaF/NaBr/NaI | 28% (w/v) PEG 4000<br>0.1 M sodium citrate<br>(pH 6.5)<br>0.1 M magnesium acetate tetrahydrate<br>0.1 M $(NH_4)_2SO_4$ |

* Values in parenthesis correspond to highest resolution shell.
[a] $R_{merge}$=Σ |I-<I>|/ΣI, where I is the observed intensity and <I> is the averaged intensity from multiple observations.
[b] <I/σI> = averaged ratio of the intensity (I) to the error of the intensity (σI).
[c] $R_{work}$=Σ|Fobs-Fcal|/Σ|Fobs|, where Fobs and Fcal are the observed and calculated structure factors, respectively.
[d] $R_{free}$ was calculated using a randomly chosen subset (5%) of the reflections not used in refinement.
[e] The original diffraction in these datasets were severely anisotropic in higher resolution and were analyzed for anisotropy and processed by STARANISO.

Table S8. Primers used for qPCR, Sequence (5’-3’)

| Gene | | Primers | Reference |
|---|---|---|---|
| HBB | F | CAG TGC AGG CTG CCT ATC | Takase et el. (2023)[1] |
| | R | ATA CTT GTG GGC CAG GGC AT | |
| HBG1/2 | F | TGG ATG ATC TCA AGG GCA C | |
| | R | TCA GTG GTA TCT GGA GGA CA | |
| BGLT3 | F | CTA TAG ATC AAG CCT GTG CCA G | Feng et el. (2022)[2] |
| | R | CAA GAG GAT AGG AGA TTC CGT G | |
| HBBP1 | F | TTA TGC TCA CGG ATG ACC TCA AAG | Ma et el. (2021)[3] |
| | R | AAC AAT CAA TAT CAC GTT GCC TAA GAG | |
| BCL11A | F | GGA GGT CAT GAT CCC CTT CT | Zheng et el. (2024)[4] |
| | R | AAC CCC AGC ACT TAA GCA AA | |
| IP08 | F | TAC TGC AAG TCA CTC TGG TTT C | Kumar et el. (2020)[5] |
| | R | ACC ACA GCC AGT GCA TAT TT | |
| IGF2BP1 | F | AGA TTG CAC CAC CCG AAA CA | Coyne S. et el. (2025)[6] |
| | R | CAC GTA TGT GGG TCT CCA GC | |
| HIC2 | F | GCT GGC TGC TGC TCA CAT G | |
| | R | TCT TGT GGG CCC GGA AGA TG | |
| LIN28B | F | CAT CTC CAT GAT AAA CCG AGA GG | Basak et el. (2020)[7] |
| | R | GTT ACC CGT ATT GAC TCA AGG C | |

Figure S1

A

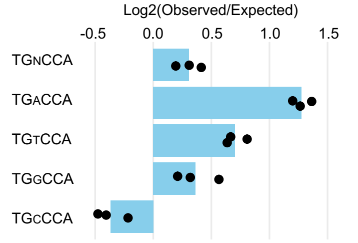


B

| Dataset | genes w/ peaks | genes with ≥1 TGNCCA | total peaks | median/ mean TGNCCA |
|---|---|---|---|---|
| Liu1 | 37 (84%) | 31 (84%) | 57 | 1 / 1.16 |
| Liu2 | 37 (84%) | 34 (92%) | 73 | 1 / 1.00 |
| Liu3 | 32 (78%) | 16 (63%) | 48 | 0 / 0.60 |

C

| Gene | Chr 11 (start – stop) | |
|---|---|---|
| *C11orf21* | 2295628 | 2303140 |
| *TSPAN32* | 2302013 | 2318204 |
| *HBBP1* | 5241954 | 5243592 |
| *HBG1* | 5248269 | 5249857 |
| *HBG2* | 5253188 | 5254781 |
| *CD82* | 44564409 | 44620358 |
| *FADS1* | 61799627 | 61817003 |
| *MIR1908* | 61815161 | 61815240 |
| | **Chr 19 (start – stop)** | |
| *GAMT* | 1397026 | 1401542 |
| *TRIP10* | 6739680 | 6751530 |
| *F2RL3* | 16888999 | 16892606 |
| *GDF15* | 18386158 | 18389176 |
| *HPN* | 35040506 | 35066573 |
| *IZUMO1* | 48740852 | 48746909 |
| *FUT1* | 48748011 | 48755358 |

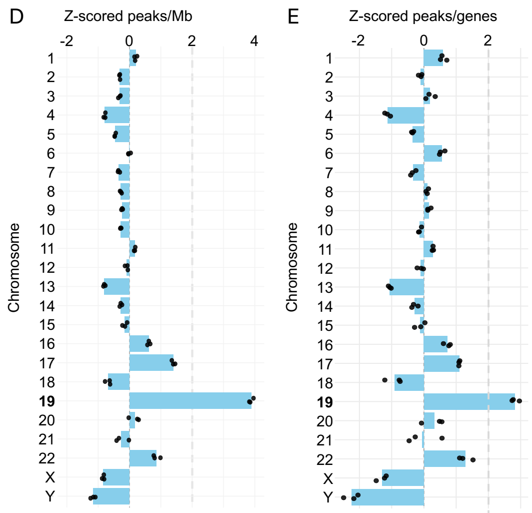


**Figure S1** (related to Figure 2). **Distribution of BCL11A peaks and TGNCCA motifs in CUT&RUN data (hg19)**. (**A**) Enrichment of TGNCCA motifs in BCL11A CUT&RUN peaks, driven primarily by higher-than-expected occurrences of TGACCA followed by TGTCCA. Expected motif frequences were estimated using first-order Markov models for each replicate. (**B**) BCL11A CUT&RUN peaks associated with BCL11A target genes that contain at least one TGNCCA motif. (**C**) BCL11A target genes located on chromosomes 11 and 19. In addition to globin genes (HBG1/2 and HBBP1), several genes have established roles in hematopoietic or erythroid biology, whereas others have no currently known functional connection. *TSPAN32* and *CD82* encode tetraspanin cell-surface proteins, with *TSPAN32* being primarily expressed in hematopoietic tissues[8,9]. The ~22-nt microRNA MIR1908 is located within an intron of the FADS1 gene, which encodes fatty acid desaturase 1 (which, in turn, controls fatty acid composition in erythrocytes[10]). *FUT1* (a fucosyltransferase expressed in hematopoietic tissues) generates the H antigen on red blood cell surfaces[11], while growth differentiation factor 15 (GDF15) in thalassemia suppresses expression of the iron regulatory protein hepcidin[12]. IZUMO1 and PUT1 (19q13.33), lie in close proximity and encode proteins with distinct biological roles (sperm-egg fusion and blood group antigen synthesis). (**D**) Z-scored BCL11A peak density normalized per megabase (Mb) or (**E**) per annotated gene, showing strong enrichment on chromosome 19. Z-scoring was computed separately for each replicate.

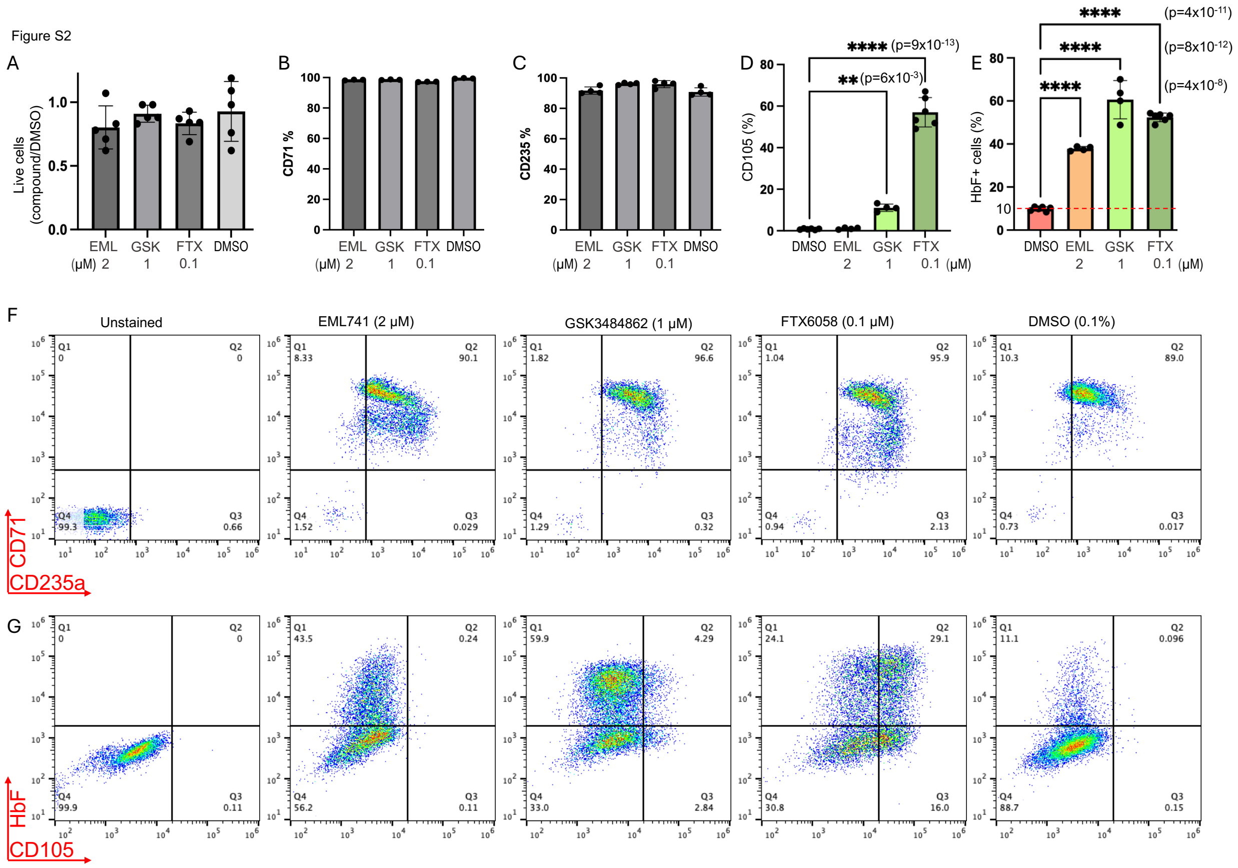


**Figure S2** (related to Figures 5 and 7). Inhibition of epigenetic silencing marks increases the population of fetal globin-expressing HUDEP-2 cells. HUDEP-2 cells were treated with three consecutive doses of each inhibitor (2 µM EML741, 1 µM GSK3484862, or 0.01 µM FTX6058) at stages 0, 1, and 2 for a total of 9 days (see Figure 5A). Stage 3 cells were harvested for analysis. (**A**) Percentage of live cells quantified using a Bio-Rad cell counter and normalized to the DMSO control. Data represent mean ± SD from N=5 independent biological replicates. (**B-D**) Flow cytometric analysis of erythroid differentiation markers CD71, CD235, and CD105. Data represent mean ± SD from N=3 or 4 biological replicates. (**E**) Percentage of fetal hemoglobin positive (HbF+) cells were determined by flow cytometry. Data represent mean ± SD from N=6 biological replicates. (**F-G**) Representative flow cytometry plot showing CD71, CD235, CD105, and HbF+ cell population in HUDEP-2 cells treated with three inhibitors, analyzed using FlowJo software.

Figure S3

A

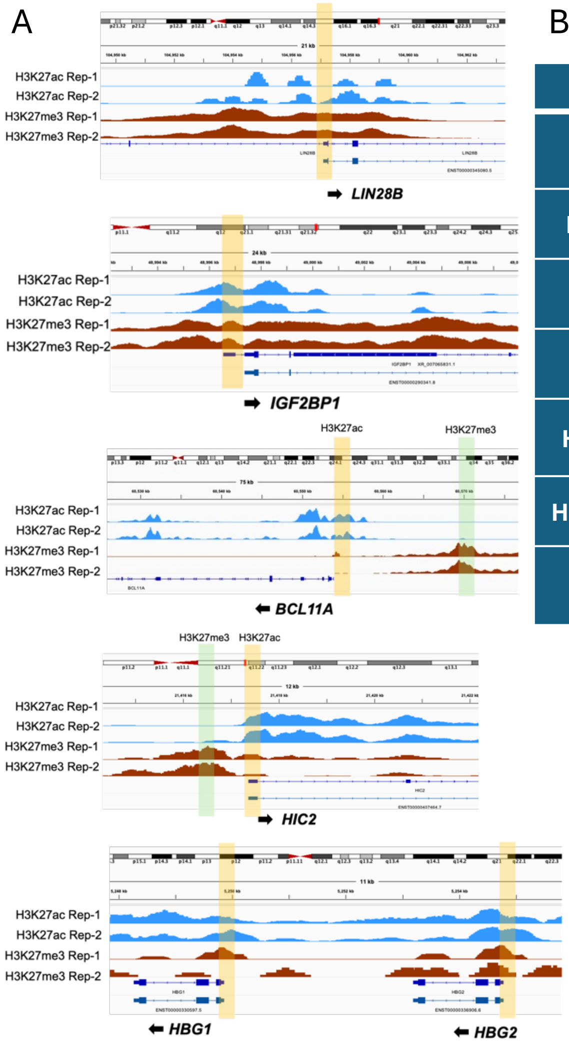


B

| Gene | | Primers (5'- 3') |
| --- | --- | --- |
| LIN28B-TSS | F | GCC AGA AAC TGG AGA GAG GA |
| | R | CGG AGT GAT CTT CTG CAT CA |
| IGF2BP1-TSS | F | AGG CTC TGG AAA CTC CAC CT |
| | R | CTA ACC GGA GCG AAC TGA AG |
| BCL11A-H3K27ac | F | CAA GCG ACT GGG TCT GAA TG |
| | R | CAG ACT CTG GCT CCT CGT TC |
| BCL11A-H3K27me3 | F | TGG GTG GTA AAG AGG CTG AC |
| | R | TAT ATT GAA ATG CGC CCA AG |
| HIC2-H3K27ac | F | AGG GCT CCC ATT ATG CAA AC |
| | R | AGG CAA GAG GAG AGG AGA CC |
| HIC2-H3K27me3 | F | TGG CGA AGG GTT ATT GAA AC |
| | R | ACT ATC GAG GTC GCC CAA GT |
| HBG1/2-TSS | F | AGC CTT GTC CTC CTC TGT GA |
| | R | GAA GCA CCC TTC AGC AGT TC |

**Figure S3** (related to Figure 6B-C). **ChIP-qPCR. (A)** Integrative Genomics Viewer (IGV) browser screenshots to show H3K27ac and H3K27me3 peaks on respective genes based on published CUT&RUN and CUT&Tag datasets available at GEO (GSE292698)[13]. The region shaded in the bar is location used for primer design for ChIP-qPCR analyses. (**B**) Primer sequences used for ChIP-qPCR.

A

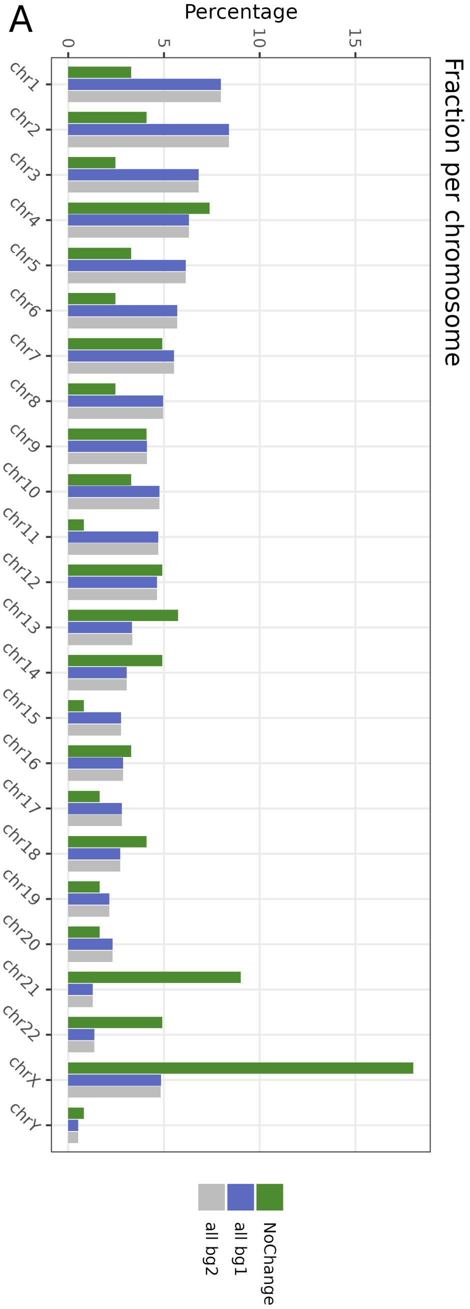


B (DMR in FTX6058-treated HUDEP-2 cells) Figure S4

| 20 hypomethylated genes | | Chromosome |
|---|---|---|
| Protein coding genes | TP63 | 3 |
| | VN1R5 | 19 |
| | ARFGAP3 | 22 |
| Pseudo genes | HNRNPKP1 | 3 |
| | RNU6-1298P | 10 |
| | BECN1P2 | 12 |
| snoRNAs | SNORA61 | 11 |
| | SNORD12 | 11 |
| Ensemble (lncRNAs or pseudo genes) | ENSG00000201245 | 1 |
| | ENSG00000228216 | 6 |
| | ENSG00000230571 | 8 |
| | ENSG00000235546 | 9 |
| | ENSG00000251517 | 11 |
| | ENSG00000254801 | 12 |
| | ENSG00000259676 | 14 |
| | ENSG00000266976 | 16 |
| | ENSG00000277967 | 19 |
| | ENSG00000279734 | 20 |
| | ENSG00000282556 | 21 |
| | ENSG00000282906 | 22 |
| **4 hypermethylated genes** | | |
| Protein coding genes | ZAR1 | 4 |
| | SLC16A6 | 17 |
| Ensemble | ENSG00000287419 | 5 |
| | ENSG00000289335 | 7 |

**Figure S4** (related to Figure 6D). DNA methylation. (**A**) The retained DNA methylation despite GSK3484862 treatment were enriched on the X-chromosome, and to a lesser extent on chromosome 21. (**B**) A small number of genes (mostly pseudogenes and non-coding RNAs) associated with differential methylated regions (DMR) following FTX6058 treatment, in comparison with DNA methylation patterns in DMSO-treated or non-differentiated HUDEP-2 cells. Among them, *TP63* level was reduced in BCL11A-KD lung squamous cell carcinoma cells[14].

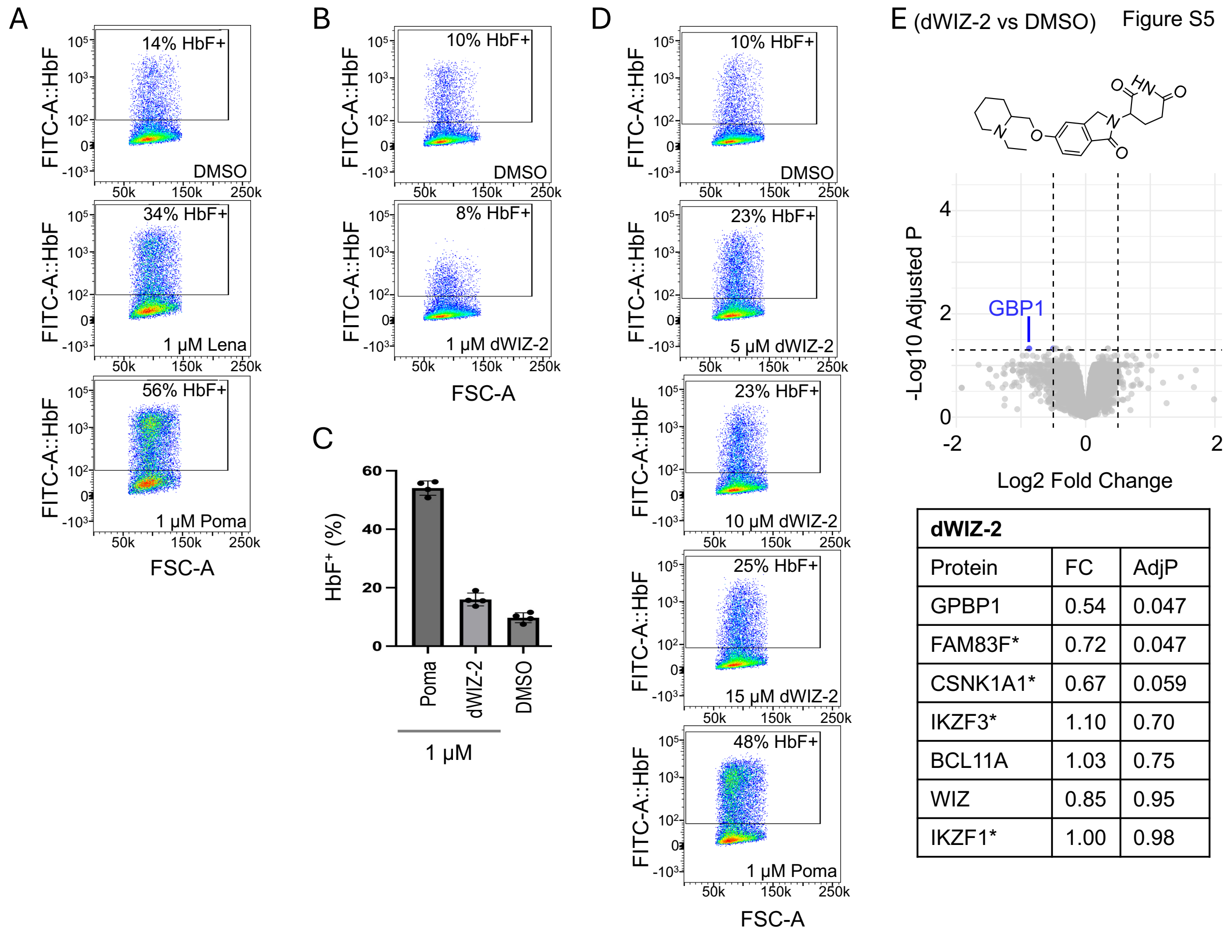


| dWIZ-2 | | |
|---|---|---|
| Protein | FC | AdjP |
| GPBP1 | 0.54 | 0.047 |
| FAM83F* | 0.72 | 0.047 |
| CSNK1A1* | 0.67 | 0.059 |
| IKZF3* | 1.10 | 0.70 |
| BCL11A | 1.03 | 0.75 |
| WIZ | 0.85 | 0.95 |
| IKZF1* | 1.00 | 0.98 |

**Figure S5** (related to Figure 8). **Commercially sourced dWIZ-2** (Selleckchem Cat.No. E4661 and batch No. E466101) **neither induces HbF induction (by flow cytometry) nor causes disappearance of WIZ (by mass spectrometry).** (**A**) Representative flow cytometry plots showing that 6 days of 1 µM lenalidomide (Lena) and pomalidomide (Poma) increases the percent of HbF-positive cells. (**B**) Representative flow cytometry plots showing that 6 days of 1 µM dWIZ-2 treatment do not increase the percent of HbF-positive cells compared to DMSO. (**C**) Bar graph of four independent replicates of 1 µM dWIZ-2 and 1 µM pomalidomide treatment following 9-day treatment. (**D**) Representative flow cytometry plots showing that 6 days of 5, 10, or 15 µM dWIZ-2 treatment increase the percent of HbF positive cells from 10% to ~22-24% compared to

48% for 1 μM pomalidomide. (**E**) Differential protein expression analysis plot of mass spectrometry data showing that WIZ is not degraded by dWIZ-2.

We also tested the recently developed HbF-inducer dWIZ-2[15], a lenalidomide derivative that decreases levels of the zinc-finger protein WIZ, a component of the histone H3 lysine 9 methyltransferase complex including at least four proteins G9a/GLP/WIZ/ZNF644[16,17]. For control, lenalidomide itself at 1 μM induced ~34% HbF (Figure S5A). Surprisingly, 1 μM of dWIZ-2 showed no induction (FC = 0.82 vs. DMSO) (Figure S5B-C), and higher doses (5-15 μM) produced only ~2-fold induction relative to DMSO, about half of pomalidomide'e effect (Figure S5D). In addition, consistent with the lack of HbF induction observed by flow cytometry, we did not detect disappearance of WIZ associated with dWIZ-2 (Figure S5E), despite using the same treatment conditions (10 μM, 6 h) reported for CD34+ hematopoietic stem cell progenitor cells[15]. We note that, beyond differences in cell type (HUDEP-2 vs. CD34+), we cannot exclude the possibility that the commercially sourced dWIZ-2 differs chemically from the compound used in the Novartis study[15].

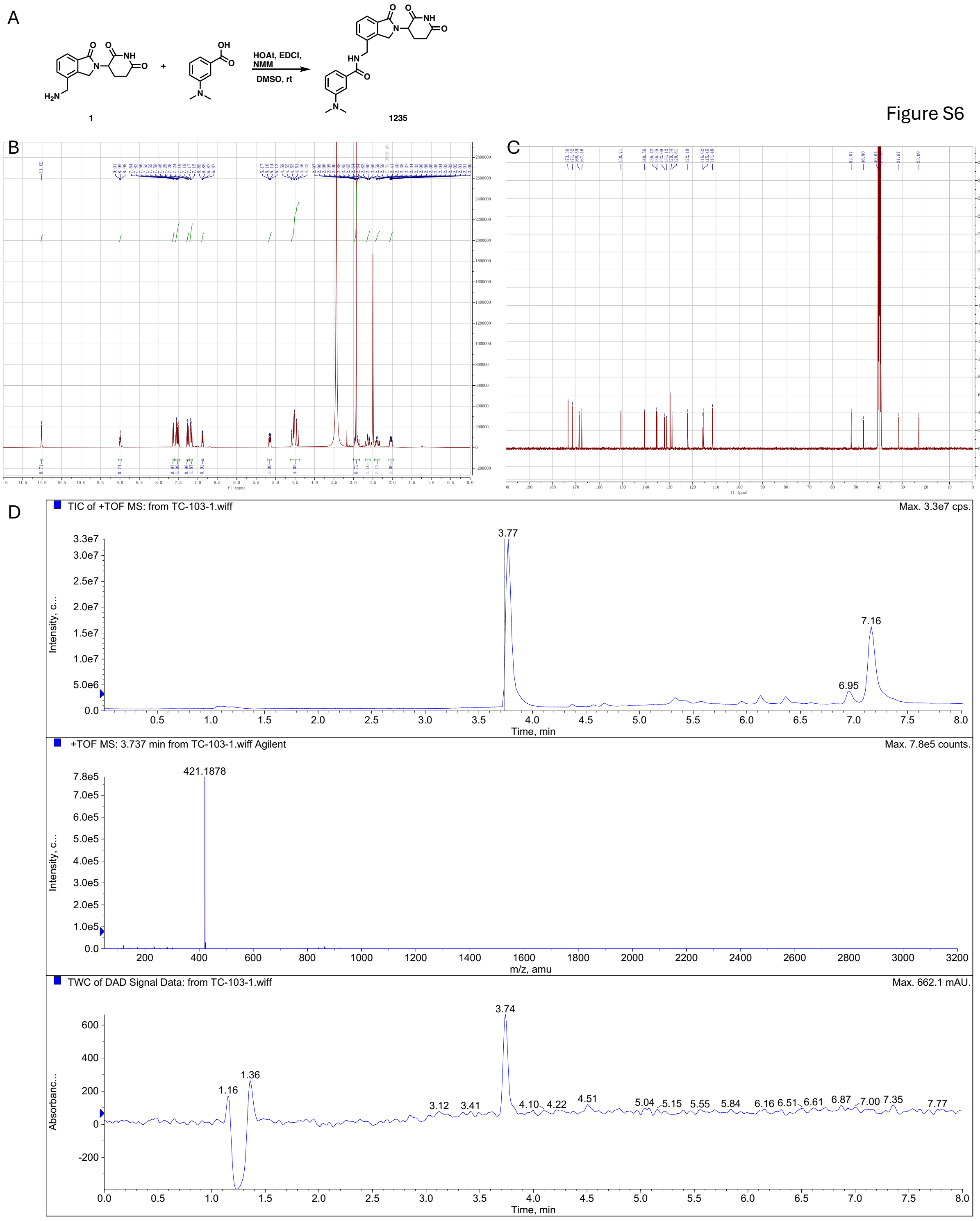


**Figure S6** (related to Figure 8G). Compound **1235**. (**A**) Synthesis scheme. (**B**) $^{1}$H NMR spectrum. (**C**) $^{13}$C NMR spectrum. (**D**) LC-MS spectra.

**References** cited in the Supplementary


1. Takase, S., Hiroyama, T., Shirai, F., Maemoto, Y., Nakata, A., Arata, M., Matsuoka, S., Sonoda, T., Niwa, H., Sato, S., et al. (2023). A specific G9a inhibitor unveils BGLT3 lncRNA as a universal mediator of chemically induced fetal globin gene expression. Nat Commun *14*, 23. 10.1038/s41467-022-35404-0.
2. Feng, R., Mayuranathan, T., Huang, P., Doerfler, P.A., Li, Y., Yao, Y., Zhang, J., Palmer, L.E., Mayberry, K., Christakopoulos, G.E., et al. (2022). Activation of gamma-globin expression by hypoxia-inducible factor 1alpha. Nature *610*, 783-790. 10.1038/s41586-022-05312-w.
3. Ma, Y., Liu, S., Gao, J., Chen, C., Zhang, X., Yuan, H., Chen, Z., Yin, X., Sun, C., Mao, Y., et al. (2021). Genome-wide analysis of pseudogenes reveals HBBP1's human-specific essentiality in erythropoiesis and implication in beta-thalassemia. Developmental cell *56*, 478-493 e411. 10.1016/j.devcel.2020.12.019.
4. Zheng, G., Yin, M., Mehta, S., Chu, I.T., Wang, S., AlShaye, A., Drainville, K., Buyanbat, A., Bienfait, F., Tenglin, K., et al. (2024). A tetramer of BCL11A is required for stable protein production and fetal hemoglobin silencing. Science *386*, 1010-1018. 10.1126/science.adp3025.
5. Kumar, J., Kaur, G., Ren, R., Lu, Y., Lin, K., Li, J., Huang, Y., Patel, A., Barton, M.C., Macfarlan, T., et al. (2020). KRAB domain of ZFP568 disrupts TRIM28-mediated abnormal interactions in cancer cells. NAR Cancer *2*, zcaa007. 10.1093/narcan/zcaa007.
6. Coyne, S., Wu, T., Hossain, M., Vinjamur, D., Starrs, M., Zeng, J., Andresen, F., Gutierrez, A., Shimamura, A., and Bauer, D.E. (2025). Direct and indirect regulation of fetal globin transcript by RNA-binding protein IGF2BP1. bioRxiv. 10.1101/2025.09.09.675143.
7. Basak, A., Munschauer, M., Lareau, C.A., Montbleau, K.E., Ulirsch, J.C., Hartigan, C.R., Schenone, M., Lian, J., Wang, Y., Huang, Y., et al. (2020). Control of human hemoglobin switching by LIN28B-mediated regulation of BCL11A translation. Nat Genet *52*, 138-145. 10.1038/s41588-019-0568-7.
8. Balise, V.D., Saito-Reis, C.A., and Gillette, J.M. (2020). Tetraspanin Scaffold Proteins Function as Key Regulators of Hematopoietic Stem Cells. Front Cell Dev Biol *8*, 598. 10.3389/fcell.2020.00598.
9. Robb, L., Tarrant, J., Groom, J., Ibrahim, M., Li, R., Borobakas, B., and Wright, M.D. (2001). Molecular characterisation of mouse and human TSSC6: evidence that TSSC6 is a genuine member of the tetraspanin superfamily and is expressed specifically in haematopoietic organs. Biochim Biophys Acta *1522*, 31-41. 10.1016/s0167-4781(01)00306-2.
10. Thomas, T., Cendali, F., Fu, X., Gamboni, F., Morrison, E.J., Beirne, J., Nemkov, T., Antonelou, M.H., Kriebardis, A., Welsby, I., et al. (2021). Fatty acid desaturase activity in mature red blood cells and implications for blood storage quality. Transfusion *61*, 1867-1883. 10.1111/trf.16402.
11. Rouquier, S., Lowe, J.B., Kelly, R.J., Fertitta, A.L., Lennon, G.G., and Giorgi, D. (1995). Molecular cloning of a human genomic region containing the H blood group alpha(1,2)fucosyltransferase gene and two H locus-related DNA restriction fragments. Isolation of a candidate for the human Secretor blood group locus. J Biol Chem *270*, 4632-4639. 10.1074/jbc.270.9.4632.

12. Tanno, T., Bhanu, N.V., Oneal, P.A., Goh, S.H., Staker, P., Lee, Y.T., Moroney, J.W., Reed, C.H., Luban, N.L., Wang, R.H., et al. (2007). High levels of GDF15 in thalassemia suppress expression of the iron regulatory protein hepcidin. Nature medicine *13*, 1096-1101. 10.1038/nm1629.
13. Wang, K., Wang, J., Feng, R., Dudnyk, K., Kim, Y.J.J., Lim, J.Y.S., Lee, M., Jr., Zhang, Y., Gao, X., Cheng, Y., et al. (2025). Silencing of BCL11A by Disrupting Enhancer-Dependent Epigenetic Insulation. Blood. 10.1182/blood.2025030211.
14. Lazarus, K.A., Hadi, F., Zambon, E., Bach, K., Santolla, M.F., Watson, J.K., Correia, L.L., Das, M., Ugur, R., Pensa, S., et al. (2018). BCL11A interacts with SOX2 to control the expression of epigenetic regulators in lung squamous carcinoma. Nat Commun *9*, 3327. 10.1038/s41467-018-05790-5.
15. Ting, P.Y., Borikar, S., Kerrigan, J.R., Thomsen, N.M., Aghania, E., Hinman, A.E., Reyes, A., Pizzato, N., Fodor, B.D., Wu, F., et al. (2024). A molecular glue degrader of the WIZ transcription factor for fetal hemoglobin induction. Science *385*, 91-99. 10.1126/science.adk6129.
16. Ueda, J., Tachibana, M., Ikura, T., and Shinkai, Y. (2006). Zinc finger protein Wiz links G9a/GLP histone methyltransferases to the co-repressor molecule CtBP. J Biol Chem *281*, 20120-20128.
17. Bian, C., Chen, Q., and Yu, X. (2015). The zinc finger proteins ZNF644 and WIZ regulate the G9a/GLP complex for gene repression. eLife *4*, e05606. 10.7554/eLife.05606.
